# Bayesian analyses of radiocarbon dates on rice reveal geographic variations in the rate of agricultural dispersal in prehistoric Japan

**DOI:** 10.1101/2022.05.25.493375

**Authors:** Enrico R. Crema, Chris J. Stevens, Shinya Shoda

## Abstract

The adoption of rice farming during the 1st millennium BC was a turning point in Japanese prehistory, defining the subsequent cultural, linguistic, genetic variation in the archipelago. Here we employ a suite of novel Bayesian techniques to estimate the regional rates of dispersal and arrival time of rice farming using radiocarbon dates on charred rice remains. Our results indicate substantial variations in the pace of agricultural adoption within the Japanese islands, hinting at the presence of a mixture of demic and cultural diffusion, geographic variations in the suitability of rice cultivation, as well as the possible role of existing social networks in facilitating or hindering the adoption of the new subsistence economy.

**Teaser:** The adoption of rice farming in prehistoric Japan was characterised by regional episodes of slowdowns and accelerations.

## Introduction

The dispersal of agriculture, its timing, speed and the mechanisms behind its spread have long been seen as one of the most important shifts that laid both the genetic, linguistic, and cultural foundations for many regions of the world (*1, 2*) Reconstructing details and variation in this process has been a key research agenda for nearly a half of century of archaeological, linguistic, and genetic research, with much emphasis dedicated on the *mode* (e.g. demic vs cultural diffusion) and *tempo* (i.e. estimates of arrival dates and the pace of the dispersal process) of this key process.

The increasing availability of large collections of radiocarbon dates has considerably improved our ability to make inferences on the *mode* and *tempo* of the dispersal of agriculture and has empirically demonstrated that such a process was geographically diverse, revealing the complex interplay ranging from the environmental suitability of farming practices to the competition between migrant farmers and incumbent communities of hunter-gatherers (*3*–*11*). However, these studies have primarily covered continental scales, employed mixed quality samples (e.g. short-lived samples of carbonised seeds vs culture chronologies), and focused primarily on the European continent. There are notable exceptions for each of these issues (*12, 13*), but to our knowledge, a comprehensive study aiming to detect variations in the *tempo* of the dispersal processes within a smaller geographical region, and exclusively based on direct dates of seeds does not exist.

The introduction of rice farming in the Japanese islands offers an important case study for understanding the interaction between early farmers and hunter-gatherers as well as a unique opportunity to contribute to the research agenda on the *mode* and *tempo* of farming dispersal. The exceptionally large number of rescue excavations during the last 40 years (*14*), and the relatively long tradition of direct dating of carbonised seeds provide unmatched quality and quantity of data, whilst both the diverse ecological settings of the Japanese islands and the varied subsistence economy of the incumbent Jomon hunter-gatherers (*15, 16*) offers a suitable scenario to investigate the extent by which these factors contributed to the pace of the dispersal process.

The arrival of rice agriculture in the Japanese archipelago is traditionally used as a defining feature that marks the end of the *Jomon* period and the beginning of the *Yayoi* period (*17*–*19*). The former is associated with a subsistence economy largely based on hunting, fishing, and gathering, with a smaller contribution of small-scale plant husbandry (*20, 21*), and covers a chronological span of over 10,000 years with its beginnings marked by the introduction of ceramic technology. The *Yayoi* period is instead associated with the introduction of a cultural package of continental origin that is archaeologically associated with paddy fields; farming tools; moated settlements; new types of pottery, dwellings, and burials; and, at a later stage, metallurgy (*18*). This continental package was brought into the Japanese islands by migrant communities from the Korean peninsula to the northern coastal area of the south-west island of Kyushu (Fig. 1) during the 1st millennium BC (*22*), and subsequently dispersed in the rest of the Japanese archipelago via a mixture of demic and cultural diffusion (*23*–*26*). This process took several centuries, effectively determining a different timing of the economic transition in different parts of the Japanese islands (*27, 28*).

**Fig. 1.**
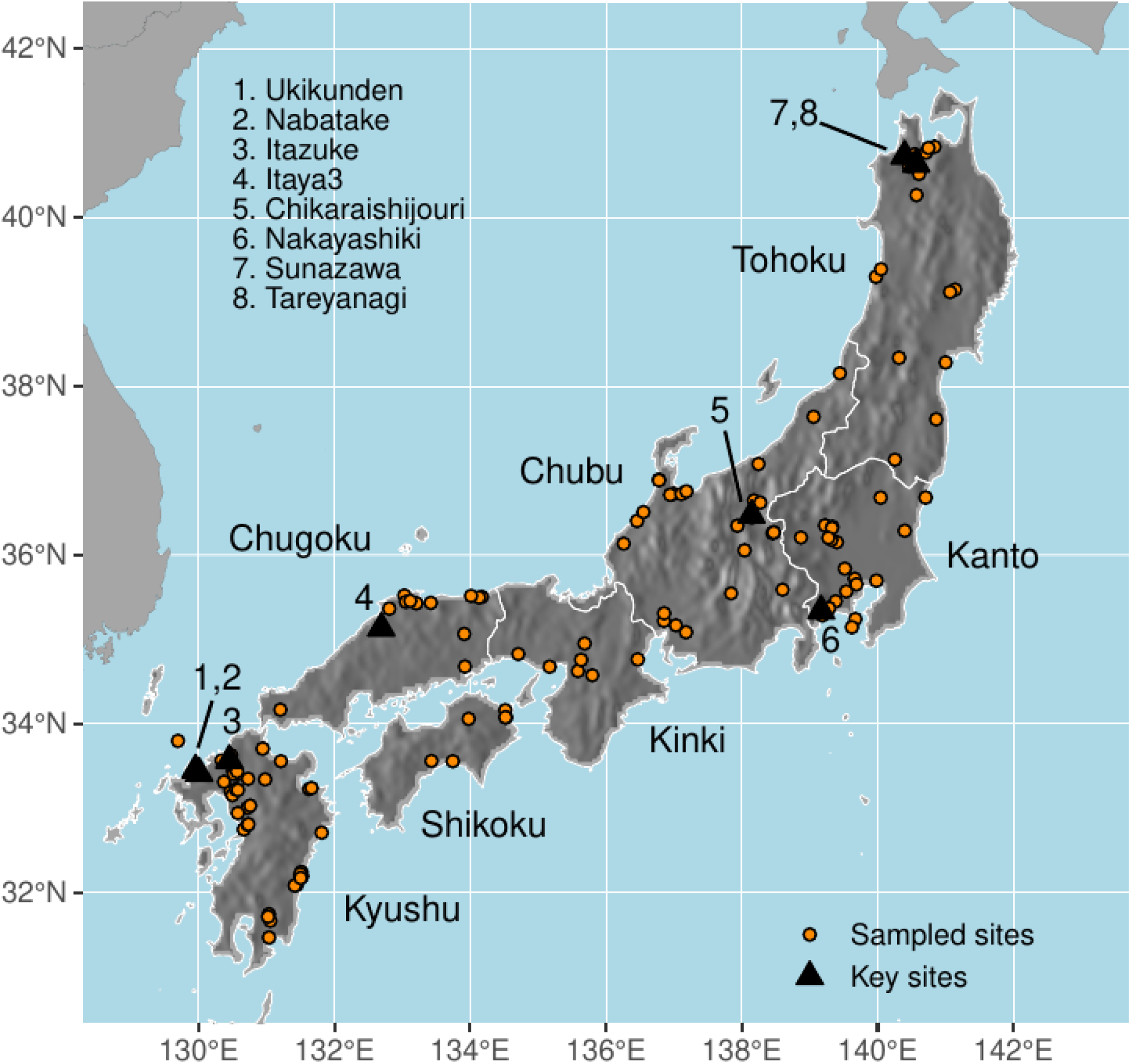
Distribution of sample Locations and key sites.

The absolute chronology of the Yayoi period has been a subject of major academic disagreement, particularly since a major breakthrough in the early 2000s (*29, 30*) where the systematic application of calibrated AMS dates shifted back its start date by nearly half a millennia. Whilst there is now a general consensus of an earlier chronology, discrepancies of a few centuries still persist amongst scholars as to when the initial establishment of rice cultivation occurred (*18, 31, 32*). The chronology of the subsequent pace of the dispersal of rice farming within the Japanese islands is even more uncertain, and has led to different suggestions on local arrival times and dispersal routes (*12, 25, 27, 31*–*33*), although there is a general consensus on a number of key features. Firstly, the dispersal of rice agriculture was limited to the three of four main islands of the Japanese archipelago (Fig. 1, Kyushu, Shikoku, and Honshu), with irrigated rice farming arriving in the Okinawa islands during the Gusuku period (AD 1187-1314, (*34*) and in the northern island of Hokkaido only during the 19th century AD (*35*). Secondly, the dispersal within the Japanese archipelago has its point of origin in Northern Kyushu, where the earliest evidence of directly dated rice grains and paddy fields are located (*31*). Thirdly, the dispersal process was characterised by several episodes of slowdowns, most notably between northern Kyushu and southern Kyushu, between Kinki and Chubu, and into northern Kanto (Fig. 1, (*25, 27*). Lastly, the dispersal of rice farming into the Tohoku region of Northern Japan (Fig. 1) showed a distinctive unusual pattern, where the earliest evidence came from the Aomori prefecture at the northern edge of the Honshu island and was possibly followed by a southward movement towards Sendai plain and subsequently to northern Kanto (*27*) where rice farming appears later than anywhere outside of Okinawa islands and Hokkaido.

Despite a general consensus on these major features of the rice farming dispersal in the Japanese islands, there are still uncertainties and disagreements on the precise chronology of arrival dates as well as the extent by which episodes of slowdowns and leapfrogs are genuinely supported by the available data and not just the result of small sample sizes. These problems are exacerbated by the constraints associated with the radiocarbon calibration plateau at around 2500 ^14^C BP, which yields calibrated dates spanning a substantial portion of the 8^th^ to 5^th^ century BC, effectively hindering accurate inferences about the timing of the dispersal within western Japan. Many of the existing studies rely on a mixture of direct dates on carbonised seeds, dates contextually and stratigraphically associated with farming (e.g. samples recovered from paddy field layers), or ceramic typology. The rich number of ceramic materials and the extremely detailed relative chronological sequences of both the Jomon and Yayoi periods have been the backbone of Japanese archaeology for decades, providing a reliable temporal reference that can be applied to any archaeological site.

The adoption of these sequences undoubtedly provides larger sample sizes and multiple proxies for the dispersal of farming that includes, amongst other things, seed impressions on pottery, paddy fields, and agricultural tools. However, the translation of these relative sequences into an absolute chronological framework has only recently been implemented, with emerging discrepancies between scholars on the start and end date of key phases (*31, 36, 37*). These dating discrepancies are exacerbated by the fact that: 1) most of the utilised radiocarbon dates are either from soot or organic residues, and hence potentially biased by old wood and reservoir effects (*38, 39*), and 2) a lack of formal statistical inferences of the start and end dates that incorporates an assessment of sampling error and calibration effects, which together contributes to chronological uncertainty (but see (*40*) for the Jomon period). As a result, there is a need to re-assess the archaeological evidence and determine where and when episodes of slowdowns in the dispersal process occurred using the most reliable samples available (i.e. direct ^14^C dates on carbonised seeds) and employing robust inferential techniques that account for the problems arising from sampling and calibration effects.

Dispersal rates of the first farming communities across the globe have typically been estimated from spatio-temporal analyses of either direct (e.g. radiocarbon dates on charred macrofossil remains, charcoal, bone collagen etc) or indirect (e.g. material culture) lines of evidence. While there is a substantial large body of literature spanning nearly four decades, with very few exceptions (*10*), the analytical framework has predominantly focused on attempts to refine the estimate of the arrival dates via Bayesian phase models (*12, 41, 42*) or on inferring the speed of crop dispersal via regression-based analyses (*3*–*6, 8, 43, 44*).

The application of Bayesian phase models for regional studies was originally introduced to study Late Glacial human occupation in NW Europe (*45*), and has since been used to study a variety of similar phenomena (e.g. (*46*) for a recent application). The key insight is to explicitly acknowledge the fact that the earliest dated evidence of farming (or any other phenomenon) simply represents a *terminus ante quem*, and hence probability distribution models are fitted to all dated samples in target regions and the ‘true’ arrival dates are obtained from the parameters of such distributions. This is effectively an adaptation of models typically designed to investigate the stratigraphic chronology of individual sites and has the benefit of taking into account the uncertainties associated with sampling and measurement errors. Furthermore, in multi-regional studies, putative expansion routes of the dispersal can be added-in as formal constraints into the model, providing an opportunity to reduce the uncertainty of the estimates. The foremost main limitation of these regional Bayesian phase models is how spatial units (i.e. ‘regions’) are being defined. Larger regions would provide more samples and lower uncertainty in the estimated parameters but at the expense of potentially missing important internal variations in arrival times. Leipe and colleagues (*12*) have recently adopted this approach to investigate the dispersal of rice farming in eastern Japan. Their work represents the first statistical assessment of the arrival time in these regions, confirming the earlier chronology of northern Tohoku compared to Southern Tohoku, and even suggesting an earlier uptake of farming in the Chubu mountains before northern Kyushu (although this latter result was entirely dependent on a single outlier date and the comparison was not made against a similar Bayesian model for Kyushu, see (*37*). However, the extent of the internal regional variation of the dispersal process is unclear and is strictly dependent on how the regional units have been defined. For example, their analysis suggests a starting date of rice farming in the Kanto region around the start of the 6th century BC, several centuries before the chronology suggested by previous authors (cf. (*27*) and an earlier date compared to their own estimate for northern Tohoku (ca. 3rd century BC). This either indicates that the region with the greatest resistance to the dispersal of farming is indeed southern Tohoku, or that such a boundary was located somewhere within the Kanto region (as suggested by other authors, e.g. (*25*). These differences in a few hundred kilometres can have profound implications in examining different hypotheses on why the dispersal of rice agriculture slowed down.

As for most common statistical models, a key assumption in the regional phase models is that dates are randomly and independently sampled. While strictly speaking these assumptions are never truly met, the problem of sample independence is particularly noteworthy when some sites offer a disproportionate number of dates. Ignoring this inconsistency is effectively the same as considering a sample of 20 dates from 20 sites for a given region to be equivalent to a sample of 20 dates from a single site from that same region. The former would be more representative of the regional arrival date for agriculture, while the latter effectively provides an estimate of the arrival date to *that particular site*. While these represent two opposite extremes, the pitfalls of not accounting for sample independence when the multiple dates are associated with the same site can potentially lead to wrong estimates (see fig. S16 and supplementary text, section 3.1 for a demonstration with a simulated dataset). The problem can be solved by either reducing the data to the earliest dated sample from each site or employing a more complex model that accounts for the hierarchical structure of the data.

In contrast to Bayesian phase models, regression-based analyses are typically employed with the objective of estimating the speed of the diffusion process rather than an accurate estimate of arrival dates. These analyses typically consist of fitting to the radiocarbon dates the geographic distance between sampling locations and a putative point of origin of the dispersal process, and the estimated slope is then used to obtain the diffusion rate. The approach has seen a number of variations, including the use of alternative distance metrics, formal comparisons of putative points of origin, and spatially explicit models that account for geographic variation in the dispersal process(*4, 6, 43, 44*). Although there is a substantial variation in the method being used (particularly with regard to how error ranges are calculated and reported), this wealth of case studies provides some benchmark estimates on the speed of the dispersal process, with figures around 0.6∼1.3 km/year for Europe (*5, 7, 47*), 2.4 ± 1.0 km/year for South Africa (*8*), and values from 0.45 to 2.88 for different parts of tropical South America (*11*).

While the possibility of direct comparison of dispersal rates makes these regression models highly appealing, particularly from the standpoint of reconstructing the generative process behind these patterns, there are a number of methodological issues related to the application of these methods. Firstly, in contrast to Bayesian phase models described above, these regression analyses commonly ignore measurement errors associated with individual dates and fit models using median calibrated dates. As noted by Riris and Silva (*48*) this approach effectively dismisses the uncertainty associated with individual dates, and hence in the best case scenario error estimates of the dispersal rates will be too low, and in the worst case, the estimate itself can be biased, particularly when the time range of analyses include plateaus in the calibration curve as is the case with the first half of the Yayoi period (see fig. S1). The second issue stems from how samples are selected for analyses in order to capture the earliest dates associated with farming. While theoretically justifiable, the practical decision is clearly dependent on the spatial scale of analyses (i.e. “earliest” where ν). Many earlier works have not provided the exact filtering protocol, although more recent works (*13, 49*) offer more explicit and reproducible criteria. A more practical solution consists of fitting a quantile regression model, where the relationship between the predictors and the dependent variable is based on specific, user-defined percentiles. Several authors (*43, 44, 48*) have fitted a quantile regression selecting extreme percentiles to effectively model the earliest date without the need of employing a filtering protocol and hence using the full sample available. Regression-based methods can also be adapted to investigate possible variations in the dispersal rate. For example, (*44*) fit a non-linear regression to account for the fact that sites closer to the source of rice agriculture were still undergoing domestication, and hence model a wave-front that accelerates as it moves away from the origin point.

Others have accounted for spatial variation using techniques such as geographically weighted regression (*4*) or employed some form of geostatistical interpolation (*6, 47*). These solutions can reveal key variations in the rates of dispersal, which in turn are interpreted as evidence of low vs high receptivity of farming practices or demic vs cultural diffusion processes. However, it is hard to discern whether observed variations in the rates of dispersal are genuine or just the consequences of calibration and measurement error (e.g. faster rates at plateaus and slower rates at steeper intervals of the calibration curve) or variation in sample structure (e.g. differences in sampling strategies and degree of sample independence).

Here we employ, for the first time, a Bayesian hierarchical gaussian process quantile regression (GPQR) which combines the principles of Bayesian phase model and quantile regression, whilst accounting for the full uncertainty of each date, sampling independence, and spatial variation in dispersal rates. To maximise the reliability of our dated samples, and avoid issues regarding old wood and reservoir effects or questionable stratigraphic associations of short-lived dates, we considered only direct 14C dates on charred rice grains (fig.1). Our approach accounts for the measurement error of individual radiocarbon dates using the same modelling protocol commonly employed in the Bayesian analyses of radiocarbon dates. We used quantile regression with the 90th and 99th percentile to specifically look at the distribution of the earliest local arrival dates and a Gaussian Process model to account for variation in the local dispersal rate. Results of the GPQR model have also informed the definition of geographic areas with internally homogenous local dispersal rates. Arrival times in these areas were inferred using a hierarchical version of the regional phase model described above. We considered an unconstrained model (model *a*) where the arrival date of each region was solely determined by the radiocarbon dates of the focal region and a partially constrained model (model *b*) which assumed a wave of advance dispersal in western Japan.

## Results

Average dispersal rates obtained from the Gaussian Process quantile regression ranged between 0.9 and 2.38 km per year (90% HPDI interval), with negligible variation between the 90th and 99th percentile models (fig. S14-15; tables S1-S2). Median posterior estimates (1.42km/year for the 99th percentile and 1.33 for the 90th percentile model) were slightly higher than the average rate of agricultural diffusion observed in Europe (*5, 7*) and the difference is even stronger in the case of a non-spatial version of the model, which yielded a median estimate of 2.00 km/year (90% HPDI interval 1.59∼2.51 km/year; fig. S3). The difference between the spatial and non-spatial model was most likely determined by the fact that the quantile regression is designed to capture the distribution of extreme observations (in this case the earliest dates), and hence biased towards the fastest estimates for a given distance. In the GPQR this is accounted for as a local deviation, whilst in the non-spatial regression, this is part of the global model.

Our analyses revealed substantial geographic variation in the dispersal rate (see Fig. 2) was comparable to those observed elsewhere on a continental scale with median estimates as high as 4km/year on one end and below 1km/year on the other. The model suggests that the rate of dispersal was initially slow within the island of Kyushu (below 1km/year), accelerated in the Chugoku and especially in the Kansai region (up to 4km/year), decreased its speed but maintained an above-average rate in the Chubu highlands (ca. 2km/year), before slowing down from the Kanto region northwards (1km/year or less). The exception within this general pattern can be captured in the 99th percentile model, where the northernmost region of the Honshu Island is associated with higher dispersal rates (ca. 2km/year) compared to the rest of eastern Japan, providing support for the presence of a leapfrog transmission to this area.

**Fig. 2.**
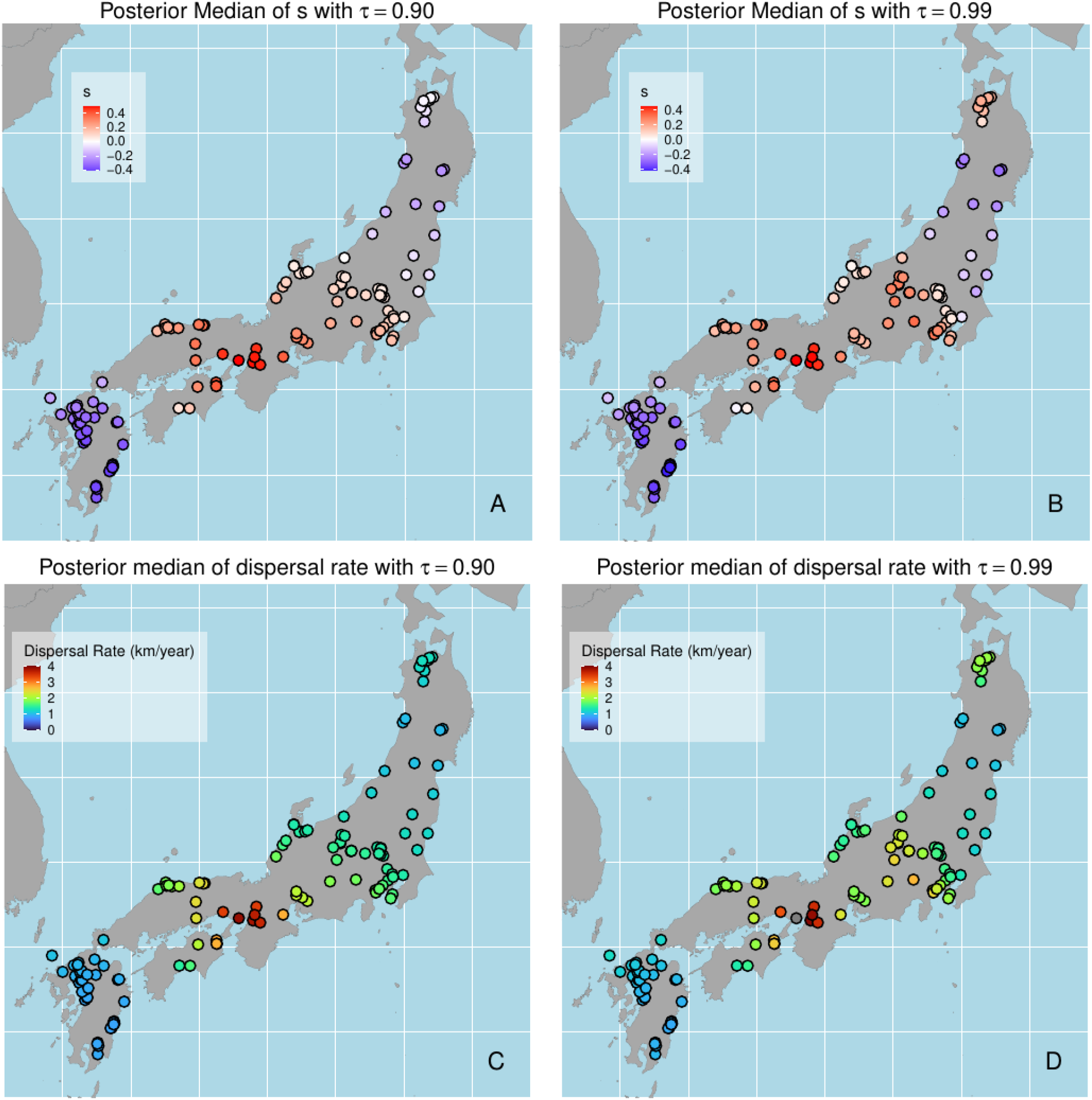
Local median posterior estimates of GPQR parameters. Deviations from the average slope parameter for the 90th percentile (**A)** and 99th percentile (**B**) models (postive values: faster dispersal rates; negative values: slower dispersal rates). Locals rates of dispersal (in km/year) for the 90th percentile (**C)** and 99th percentile (**D**) models.

Estimates of the arrival time in the different areas obtained from the Bayesian hierarchical model are aligned with these findings (Fig. 3, see also Fig S18-19 and table S4). In the unconstrained model (model *a*), the estimated arrival date of rice farming in northern Kyushu (Area I) is between 1176 and 845 BC (90% HPDI, see table S5), confirming this area to be that with the earliest adoption of rice farming in Japan (*contra* (*12*), see fig. S20). The next two earliest areas, comprising Chugoku, Shikoku and Kansai, have estimated arrival dates between 1061 and 779 BC for Area III (90% HPDI) and between 946 and 703 BC for Area IV (90% HPDI) in the constrained models, and with wider posterior density intervals in the unconstrained models. Arrival dates for Area II (central and southern Kyushu) and V (Chubu mountains) are very similar, the former yielding estimates between 735 to 430 BC (90% HPDI) and the latter between 754 to 560 BC (90% HPDI) in the constrained model. The largest chronological gap between geographically adjacent regions is recorded northeast of Area V, the same regions where we start to observe below-average local rates of dispersal in the GPQR model. Estimates of arrival time in Area VI (most of Kanto region, excluding Kanagawa prefecture) is between 471 and 124 BC, ca. 375 years (90% HPDI: 133 ∼ 557 years, fig. S21) after the arrival in Area V, whilst Area VII yielded the latest median arrival date (152 BC, 90% HPDI: 434BC to 42 AD) in the constrained model. As for the results of the GPQR with the 99th percentile, the phase model does confirm that rice cultivation arrived in northern Tohoku (Area VIII) before neighbouring regions (Areas VI and VII), with an estimated arrival time between 709 and 203 BC (90% HPDI) in the constrained model. While the comparatively smaller sample size for this region has led to fairly large posterior intervals, the unconstrained model nonetheless provides support for a leapfrog transmission in northern Tohoku, with the probability of rice arriving in areas VI and VII before VIII being respectively 0.19 and 0.12 (fig. S20).

**Fig. 3.**
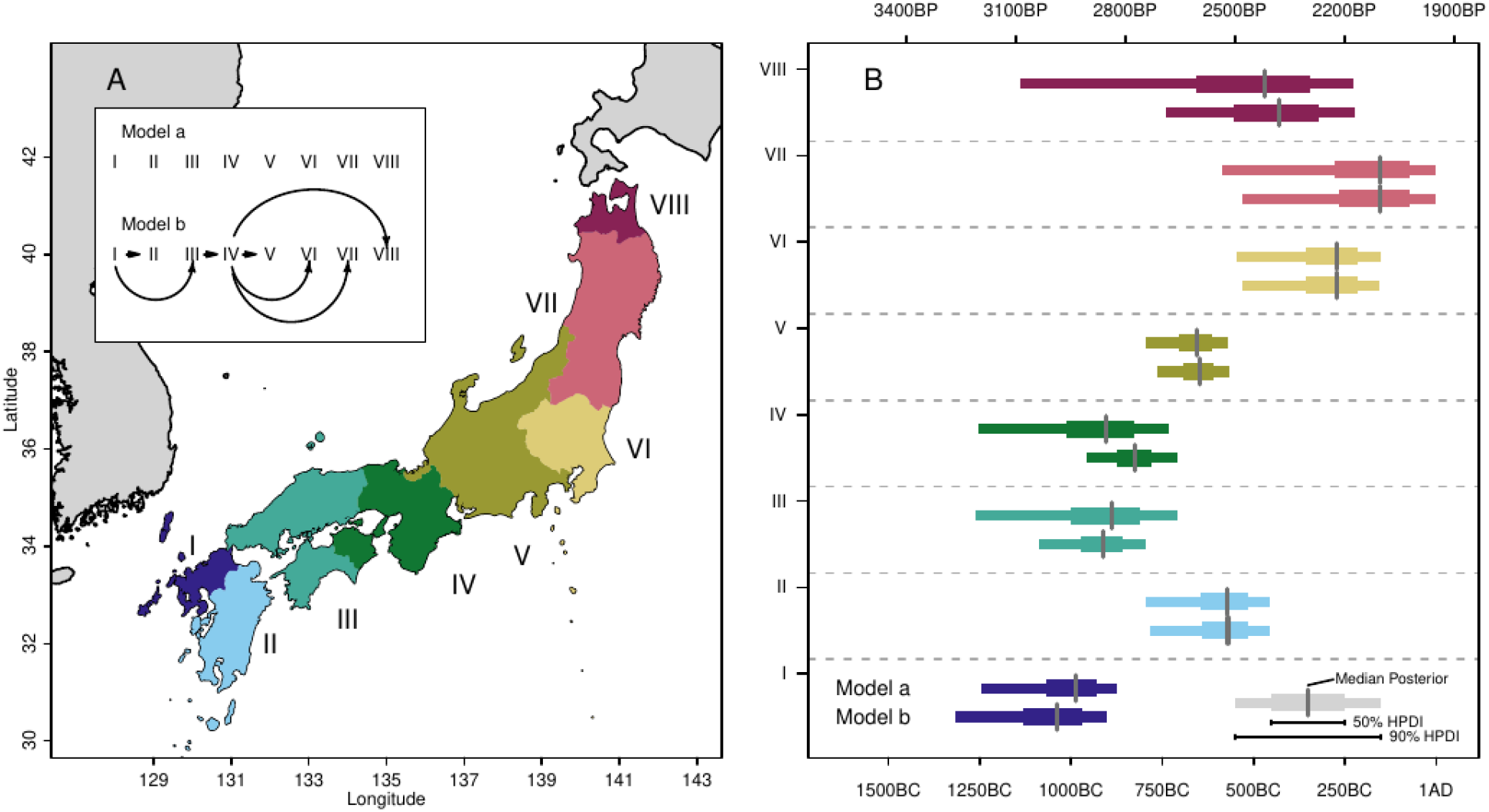
Estimated arrival dates of rice in different areas. Geographic areas used for the analyses and the constraint relationships used in the two hierarchical models (**A**). Posterior distribution of ν_k_ (i.e. arrival date) for the two models in the eight areas analyses.

## Discussion

The dispersal of rice farming in the Japanese islands during the 1st millennium BC represents a defining moment that sets the foundations of present-day genetic, linguistic, and cultural variations in the archipelago (*50*–*52*). The Bayesian analyses of charred rice dates suggest that this process was characterised by substantial regional variation in its pace, with local episodes of increase and decrease speed in the rate of dispersal.

From a methodological standpoint, our analyses address many of the issues affecting previous studies attempting to estimate rates of dispersal and arrival times. We demonstrated that inferring front speeds ignoring the uncertainty associated with individual dates can drastically change estimates, particularly in the presence of calibration plateaus (see fig. S1 and S2) and that ignoring sample independence can impact the conclusion of phase models applied in regional settings. Our Gaussian process model also provides an alternative to simple geostatistical interpolations of arrival dates employed in previous studies and offers a robust method for detecting episodes of slowdowns in the dispersal rates. Although the relative density of dates on short-lived samples is higher compared to studies with continental scales of analyses, our limited sample size led to fairly larger posterior estimates. This is, however, in part due to the fact that our analyses fully account for the measurement errors associated with individual dates in contrast to previous studies based on median dates. The choice of limiting the sample to direct dates on charred rice, while overcoming issues of contaminated dates, does exclude other potentially more reliable proxies of farming, such as paddy field sites or the presence of dedicated cultivation and harvesting tools, as well as the possibility that in some cases charred remains of rice might have resulted through trade and exchange rather than local production.

Nonetheless, our findings provide a formal assessment of existing dispersal models in the literature and whilst it has confirmed some of the key narratives that have been proposed, it offers new insights as well as a more robust chronological framework. In general terms, our estimates point to earlier arrival dates for all regions, as, in contrast to most existing studies, we base our figures on formal inference and not on the visual inspection of sample dates (but see Leipe et al 2020 for an exception). Our model suggests that the introduction of rice agriculture in northern Kyushu occurred at the turn of the 1st millennium BC. Evidence for rice farming occurring in the Japanese islands during this time or earlier is patchy and often controversial, though their density is higher in northern Kyushu compared to other parts of the archipelago (cf. fig. 83, (*33*). Several studies have shown impressions of rice grains on potsherds pre-dating the Initial Yayoi period (*37*). Nakazawa and Ushino (*53*) identified a rice grain impression from a ceramic vessel recovered at Itaya III site in Shimane prefecture (Fig. 1) attributed to the Maeike phase of the Final Jomon period. However, its typological classification is controversial (see (*54*) for suggestions of a later chronology) and the lack of direct dating and a reliable absolute chronology of the Maeike phase does not provide sufficiently robust evidence for farming occurring earlier in the Chugoku region. Leipe and colleagues (*12*) argue for the possibility of an earlier adoption of rice farming in the Chubu highlands based on a single direct date of single rice grain (IAAA-83092, 2889±29 ^14^C BP) from the Chikaraishijori site in Chubu (Fig. 1). However, the sample is a clear outlier when compared to other charred rice dates from the same site, and one of the authors acknowledges in a subsequent study that its’ identity could not be verified (*37*). Removing this sample provides a much later date for this region, and our model (fig. S20) strongly indicates that the arrival of rice in northern Kyushu (Area I) predates the arrival in the Chubu Highlands (Area V).

The analyses show divergent speeds for the expansion of agriculture outside northern Kyushu. The eastward expansion within the cultural area of Tottaimon pottery (Areas III & IV) was fast-paced, reaching dispersal rates of over 4km per year, whilst the adoption of farming in southern Kyushu was significantly slower with later arrival dates compared to Chugoku, Shikoku and Kinki regions (figs.2 & S20). The fast dispersal rate in Chugoku, Shikoku, and Kinki has been pointed out by several authors in the past (e.g. (*27*)), although our chronology is substantially earlier, particularly for Kinki (Area IV). The slower dispersal rate in the rest of the island of Kyushu is less discussed in the literature (but see (*55*). Evidence of earlier farming sites in areas south of northern Kyushu has been advocated, but most are based on indirect lines of evidence and a reliable chronological assessment is still lacking (*33, 34*).

The expansion of rice farming in central and eastern Japan is characterised by a slower pace compared to that seen for Areas III and IV albeit with considerable variability. The adoption of early wet-rice farming in central Japan is noteworthy, mostly due to the lack of paddy field sites dated to this period in the area, which has led some authors to suggest the presence of dry-rice farming e.g.((*25*) for Nakayashiki site in Kanagawa, Fig. 1; but see (*56*) for an alternative interpretation). While the pace of dispersal is slower than in Kinki, the presence of hybrid pottery styles (i.e.. Joukonmon pottery) does hint at the presence of different forms of cultural interaction (*57*), although our analyses suggest that the front speed in this area was faster than previously thought. Kobayashi (*25*) identifies two “Jomon Walls” in the area, one located near the waist of Japan (*57*), approximately between Area IV and Area V (the “Chubu Wall”), and one located in the Kanto region (the “Tone River Wall”, between Areas V and VI). The former “wall” corresponds to the expansion limit of the Ongagawa pottery (*58*), and has been considered by many as the point of the largest slow-down in Honshu. Our analyses confirm a slower rate of dispersal in eastern Japan compared to most of western Japan, but we identify that the largest discrepancy in arrival dates appears to be between Areas V and VI (Fig. 3 and fig. S21).

The expansion of rice farming beyond Area V is further characterised by higher levels of uncertainty, particularly due to the smaller sample sizes in the Tohoku region. Our analyses do confirm that rice farming was present in the Tsugaru plain in Aomori prefecture (Area VIII) before other regions in eastern Japan (Areas VI and VII, fig. S20). However, we found no evidence supporting a coastal route via the sea of Japan followed by a possible downward expansion via the Pacific coast as claimed by some authors (*27*) based on the presumed movement of material culture (*59*). Instead, our analyses identified southern Tohoku (Area VII) to be the last region of those analysed to witness the arrival of rice farming (although with considerable uncertainty, see fig. S20).

Observed variations in the pace of rice farming dispersal can potentially be explained by possible differences in the mode of transmission (i.e. demic vs cultural), the density of incumbent populations of hunter-gatherers and their transmission network, and variation in the suitability of rice cultivation due to local topographic settings and climatic conditions. At the turn of the 1st millennium BC, the Japanese archipelago was characterised by three major ceramic zones (Ongagawa in the west, Fusenmon in the centre, and Kamegaoka in the northeast; (*18, 58*) and higher population densities in eastern Japan compared to the west (*60*). With other things being equal, this setting could explain the fundamental east-west divide in the dispersal rate. In western Japan, expanding migrant communities potentially had less competition for space and at the same time might have become integrated into pre-existing social and inter-marriage networks. The slight slow-down in central Japan can then be attributed to the consequence of crossing a cultural boundary that brought expanding agriculturalists into contact with communities outside the Tottaimon social network, along with a necessary adaptation to the different topographic settings of central Japan, characterised by smaller coastal plains and narrower fluvial valleys. The slower dispersal rate of rice farming in the Kanto and Tohoku regions can then be ascribed to different processes, in which elements of cultural rather than demic diffusion became the dominant driving forces compared to western regions of the archipelago (see also Hayashi 1985). The stronger presence of incumbent material culture traits in eastern Japan during the Yayoi period seems to support this hypothesis, although it is worth highlighting that a pure cultural transmission of a complex technology such as rice farming would have been nearly impossible. Despite this it is worth noting that early paddy field sites, such as Sunazawa and Tareyanagi in Aomori prefecture (Area I), were not associated with the same farming toolkits found in other paddy field sites, hinting at the possibility of a different transmission process. These early paddy field sites were, however, abandoned after just a few centuries, and the local communities reverted to a predominantly hunting and gathering based economy, suggesting that rice farming was not a fully consolidated part of their subsistence economy. Takase (*61*) suggests these abandonments were triggered by a local flood event, but the lack of subsequent recovery of rice agriculture hints at the possibility that northern Tohoku was at the edge of the thermal niche for rice cultivation.

The picture that emerges from these analyses supports the existence of a variety of different transmission mechanisms that had been previously implied by other lines of archaeological evidence. The heterogeneous ecological settings and the different regional trajectories taken by the incumbent Jomon population have effectively set a complex mosaic within which the dispersal of rice agriculture took place. The extent to which different social, cultural, and environmental and environmental contexts acted as friction to the dispersal itself is outside the scope of this paper, but the methodological framework introduced here can set the grounds for inferring these.

## Materials and Methods

### Radiocarbon data

We compiled a ^14^C dataset of 439 charred samples of direct dates on *Oryza sativa* grains from 218 archaeological sites located in the Japanese islands. Dates were collected from site reports, journal articles, and the ^14^C database of the National Museum of Japanese History (*62*). We filtered this data by excluding samples from the Ryukyu Islands and Hokkaido, samples yielding uncalibrated ^14^C ages smaller than 1,000 bp, and those with possible contamination of dated carbon or without a reliable taxonomic identification. The resulting dataset (data file S1), consisted of 294 dates from 132 site locations.

### Gaussian Process Quantile Regression

We modelled the geographic variation in the rate of dispersal from a putative origin point located at Ukikunden Shell Midden in Northern Kyushu, where the earliest charred rice date in our dataset was recovered, using a Bayesian hierarchical Gaussian process quantile regression (GPQR) defined as follows:

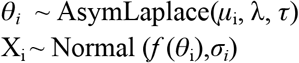

where *θ*_*i*_ is the true calendar date of the earliest charred sample identified at each site *i, f* (*θ*_i_) is its corresponding ^14^C age on the IntCal20 calibration curve (*63*), *X*_*i*_ is the observed conventional radiocarbon age of the sample, and *σ*_*i*_ is the square root of the sum of the squares of the samples’ ^14^C age error and the error on the calibration curve. The core of the model is the asymmetric Laplace likelihood (*64*), where *τ* is the quantile of interest, λ is the scale, and *μ*_i_ is the location parameter defined by the linear expression:

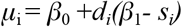

where *β*_0_ is the intercept, *d*_*i*_ is the great-arc distance (in km) between the focal site *i* and the putative origin point, *β*_1_ is the average negative reciprocal of the dispersal rate, and *s*_*i*_ is a spatially autocorrelated random effect representing the local deviation of the dispersal rate. More specifically, *s*_*i*_ is modelled as a multivariate normal distribution:

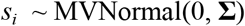

with a vector of mean equal to 0 and the covariance matrix defined by a quadratic exponential model:

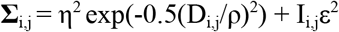

where the covariance **Σ**_i,j_ between pairs of sites *i* and *j* declines exponentially as a function of their great-arc distance *D*_i,j_ at a rate defined by the length scale parameter ρ, and with a maximum covariance equal to the square of the marginal deviation η. The term I_i,j_ ε^2^ provides additional covariance ε^2^ in case the *i* and *j* are identical (i.e. I_i,j_ is an identity matrix equal to 0 when *i* ≠ *j* and 1 when *i = j*). In practical terms, because there is only one local deviation per site this part of the equation is not relevant, however with small values of ρ and *D*_i,j_ setting ε^2^ to 0 can be problematic for algorithmic reasons (non-positive eigenvectors), and hence a small constant of 10^-6^ was assigned to ε^2^. We also considered a non-spatial version of the same model where *μ*_i_ was simply defined by a linear equation without a random effect, i.e. *β*_0_ +*d*_*i*_*β*_1_ (see supplementary text, section 1.1).

We used the weakly informative priors informed by prior predictive checks and realistic ranges of dispersal rates inferred from other studies (see supplementary text, section 2.1, fig. S7-S8). To establish the robustness of the proposed approach, we generated a simulated dataset with a comparable sample size to our observed data and fitted our GPQR model (see supplementary text, section 2.2, fig.S9-S11). Results indicate a good performance with all fixed parameters and the majority of random effect parameters within the 95% higher posterior density interval.

Following previous studies, we fitted our models using the 90th and the 99th percentile (i.e. *τ*=0.9 and *τ*=0.99). The latter represents more closely the earliest arrival date of rice but it is more susceptible to potential outlier dates.

### Hierarchical Phase Model

We estimated the arrival date for eight geographic areas using a Bayesian hierarchical phase model (see supplementary text, section 3). The spatial extent of the areas was defined based on a combination of prior archaeological knowledge whilst ensuring that the dispersal rates estimated by the GPQR were internally homogenous. The eight areas are: I (Fukuoka, Saga, and Nagasaki prefectures); II (Oita, Miyazaki, Kagoshima, and Kumamoto prefectures); III (Chugoku region, Ehime, and Kochi prefectures); IV (Kansai region, Kagawa, and Tokushima prefectures); V (Chubu region and Kanagawa prefecture); VI (Saitama, Tokyo, Chiba, Gunma, Tochigi, and Ibaraki prefectures); VII (Fukushima, Yamagata, Miyagi, Iwate, and Akita prefectures); and VIII (Aomori) (table S3). The separation of the island of Kyushu between Areas I and II was based on the distinct nature of the earliest farming communities of the northern Kyushu, while eastern Shikoku was assigned to Area IV rather than III based on the result of the GPQR that suggested a faster dispersal rate in those areas closer to sites in eastern Kansai. Kanagawa had several early sites and faster dispersal rates than the rest of Kanto and hence was assigned to Area V, and finally, Aomori was kept separated from the rest of Tohoku (Area VII) to evaluate the leapfrogging hypotheses based on the presence of earlier paddy fields in the Hirosaki plain area.

In contrast to typical regional phase models where sample independence are either ignored (i.e. multiple dates from the same site represented within each phase) or controlled by limiting the number of specimens to a unit per site, we first modelled the distribution of all charred rice dates within each site *i* and estimated its start date ν_*i*_ as follows:

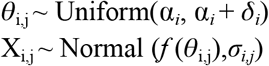

where *θ*_i,j_ is the calendar date of the sample *j* from the site *i*, and *δ*_*i*_ is the duration of the rice-use at the focal site. The measurement error of the *θ*_i,j_ was modelled using the same procedure used in the GPQR model. The distribution of start parameters ν_i_ within a given Area k was modelled using a uniform probability distribution with start and end dates ν_k_ and ν_k_, where the former is our primary parameter of interest representing the arrival of rice in the focal region. As for the GPQR model above, we tested the robustness of our model on simulated data (see supplementary text, section 3.1).

We considered two different models based on assumptions (or lack thereof) on the relationship of ν_k_ for the eight areas (see Fig. 3, left panel). In **model a**, we assumed no constraint, and hence estimates of ν_k_ were effectively made independently for each area. In **model b**, we assumed instead a wave of advance dispersal between northern Kyushu (Area I) and Kansai (Area IV), hence imposing the constraints ν_I_ > ν_II_ and ν_I_ > ν_III_ > ν_IV_, so that Area I was earlier than Area II, Area III was earlier than Area IV etc. Areas in central and eastern Japan (Areas V∼VIII) were assumed to be later than Area IV, but we did not impose any constraints between them (i.e. ν_IV_ > ν_V_, ν_IV_ > ν_VI_, ν_IV_ > ν_VII_, and ν_IV_ > ν_VII_) to allow for possible leapfrog dispersals as hypothesised by some authors (e.g. (*27*). We used flat priors bounded between 5000 and 500 cal BP for ν_k_ and ν_k_, and weakly informative prior for *δ* (see supplementary text, section 3.2) for both models.

### Parameter Inference

Model fitting was carried out in R v.4.1.0 (R Core Team 2021), using the *nimble* v. 0.12.1(*65, 66*) and the *nimbleCarbon* v.0.2.0 (*67, 68*) R packages. We used a Metropolis-Hasting adaptive random walk sampler for all parameters with the exception of *β*_1_ and *s*_*j*_ in the GPQR model, which were inferred using an automated factor slice sampler to account for correlation between the parameters. We ran four chains for all models, using 2 million iterations for the GPQR and 6 million iterations for the hierarchical phase model. In both cases, we dedicated half the iterations for the burn-in and thinned our sample to reduce file sizes (every 300 steps in the phase model and 100 in the GPQR). Convergence of the chains was checked using the Gelman-Rubin diagnostic and visual inspection of the trace plots.

## Acknowledgements

We would like to thank Yuichiro Kudo for providing additional details from his radiocarbon database, Kie Yi for supporting the additional data collection and the ENCOUNTER team (Leah Brainerd, Simon Carrignon, and Oliver Craig) for the continuous support.

## Author contributions

Conceptualization: ERC, CJS Methodology: ERC Statistical Analyses: ERC

Data Collection: ERC, CJS, SS Writing—original draft: ERC, CJS Writing—review & editing: ERC, CJS, SS

## Data and materials availability

All data and code necessary to reproduce the analyses are available at the github repository https://github.com/ercrema/yayoi_rice_dispersal and permanently archived on <URL to be provided after acceptance>..

## Supplementary Materials Text S1

**Text S1**.Supplementary text.

**Figure S1**. Impact of systematic measurement error on a putative relationship between distance and time for dates falling the calibration plateau. True rate (black) shows a positive slope indicative of a dispersal rate of ca 2.85 km/year, whilst median calibrated dates (red) have a flat slope with an instantaneous (i.e. infinite) dispersal rate.

**Figure S2**. Quantile regression model fitted on median calibrated date (blue line) and accounting for measurement error via Bayesian hierarchical model (red line). Hollow dots are the median calibrated dates while the filled dots are the median posterior of the Bayesian model accounting for the overal model. The offset between median dates and the model posterior are the largest in the calibration plateau region, where the calibrated dates generally show flat distributions and the overall model can provide the highest amount of additional information.

**Figure S3**. Estimated dispersal rate for the spread of charred rice farming in Japan using Bayesian quantile regression.

**Figure S4**. The quadratic exponential model and its relationship to its parameters ρ, η, and d_i,j_ assuming *i≠j*.

**Figure S5**. Simulated local dispersal rates with different settings of ρ and η, with 300 site locations and *β*_0_ = 3000 and *β*_1_ = 1.

**Figure S6**. Relationship between regression slope (here *s-β*_*1*_) and its negative reciprocal, i.e. rate of dispersal.

**Figure S7**. Prior predictive check for β_0_ (modelled as a Gaussian with a mean of 3000 and a standard deviation of 200), β_1_ (modelled as an exponential with a rate of 1), and η^2^ (modelled as an exponential with a rate of 20). The slope was derived by sampled values of s - β_1_, with s drawn from a gaussian with a mean of 0 and a standard deviation equivalent to η. Dashed lines represent the slope for specific rates of dispersal.

**Figure S8**. Prior predictive check for η^2^ (modelled as an exponential with a rate of 20) and ν (modelled as a truncated gamma distribution with a shape of 10 and a rate of 0.06, bounded between 1 and 1350).

**Figure S9**. Simulated and predicted (median posterior) dispersal rates.

**Figure S10**. Simulated and predicted values of the spatial random effect parameter *s*.

**Figure S11**. Posterior distribution of key model parameters for the simulated dataset with corresponding “true” values shown as a dashed line. Notice that for *β*_0_ the line represents the 90th percentile matching the value of *τ*used for inference.

**Figure S12**. Trace plots for *β*_0_, *β*_1_, ρ, and η^2^ for the *τ*=0.9 GPQR model.

**Figure S13**. Trace plots for *β*_0_, *β*_1_, ρ, and η^2^ for the *τ*=0.99 GPQR model.

**Figure S14**. Marginal posterior distribution of *β*_0_, 1/*β*_1_, ρ, and η^2^ in the *τ*=0.90 GPQR model.

**Figure S15**. Marginal posterior distribution of *β*_0_, 1/*β*_1_, ρ, and η^2^ in the *τ*=0.99 GPQR model.

**Figure S16**. Marginal posterior distribution of the start and end dates ν and ν in the hierarchical and non-hierarchical models .

**Figure S17**. Prior predictive check for the duration parameter δ. Generating using 5,000 random samples from relevant priors.

**Figure S18**. Marginal posterior distribution of ν for model *a*.

**Figure S19**. Marginal posterior distribution of ν for model *b*.

**Figure S20**. Probability of row region’s ν being after column region’s ν for model *a*

**Figure S21**. Posterior estimates of pairwise difference (column minus row) of ν in model *a*. Highlighted regions represent the 90% HPDI (red: row is earlier; blue: row is later).

**Table S1**. Summary statistics and diagnostics for *β*_0_, 1/*β*_1_, ρ, and η^2^ in the *τ*=0.90 GPQR model.

**Table S2**. Summary statistics and diagnostics for *β*_0_, 1/*β*_1_, ρ, and η^2^ in the *τ*=0.99 GPQR model.

**Table S3**. Number of sites and radiocarbon dates from each area.

**Table S4**. Summary statistics and diagnostics for the ν parameters in the two models.

**Data File S1**. ^14^C Dates used for the analyses.

