## Supplementary Text, Figures, and Tables for "Bayesian analyses of radiocarbon dates on rice reveal geographic variations in the rate of agricultural dispersal in prehistoric Japan"

**for**

### 1. Handling the Impact of Radiocarbon Measurement Error and Calibration Plateau

Estimates of farming dispersal rate are typically carried out by regressing the distance of focal sites against the mean or the median calibrated radiocarbon date of the earliest evidence of farming on those sites (Pinhasi et al 2005, Van Etten and Hijmans 2010, Silva and Steele 2014, Jerardino et al 2014, de Souza et al 2020). The rate is then computed by taking the negative (in case time is represented by BP and positive values) reciprocal of the slope coefficient (but see Pinhasi et al 2005 for fitting time against distance instead). Confidence intervals of the derived rate estimates do not however account for the impact of the measurement error of individual dates, nor the fact that these errors are not random but impact systematically by calibration effects. Figure S1 illustrates this point, showing a hypothetical dispersal of 2.85 km/year with four observation points in calendar time (black points). The red dots show the corresponding median calibrated dates, which due to the effect of the calibration plateau converges to almost identical values, biassing the slope to 0, and hence an estimated instantaneous dispersal.

A possible solution, first introduced by Steele (2010), and later adopted in some studies (e.g. de Souza et al 2020, Riris and Silva 2021), consists of fitting the regression model to iteratively sampled calendar dates from each calibrated distribution and obtaining a bootstrapped estimate of the confidence interval. While this solution does account for the uncertainty of each measurement, it holds a conservative stance where effectively each point is treated separately, without acknowledging the fact that these uncertainties are potentially correlated to each other. Furthermore, these approaches often consist of aggregating the estimates of individual iterations but not their associated errors (but see Riris and Silva 2021), and hence the resulting confidence intervals could potentially fail to properly account for the uncertainty derived from sampling error.

#### 1.1 Bayesian Quantile Regression

An alternative approach is to fit a hierarchical measurement error model, following the same principles adopted in Bayesian phase models (Buck et al 1992), and more recently in radiocarbon based demographic inference (Crema and Shoda 2021). In this case, the regression model can be conceptualised as follow:

$$\theta_i \sim \text{Normal}(\mu_i, \varepsilon) \quad [1]$$

$$\mu_i = \beta_0 + \beta_1 d_i \quad [2]$$

$$X_i \sim \text{Normal}(f(\theta_i), \sigma_i) \quad [3]$$

where $\theta_i$ is the true calendar date of the date at each site focal site $i$, $d_i$ is the distance of the site from the putative source of dispersal, $f(\theta_i)$ is its corresponding $^{14}C$ age on the IntCal20 calibration curve (Reimer et al 2020), and $X_i$ is the observed conventional radiocarbon age of the sample, and $\sigma_i$ is the square root of the sum of the squares of the samples' $^{14}C$ age error and the error on the calibration curve. Equation [3] effectively handles the measurement error of $\theta$ and establishes its relationship to the observed $^{14}C$ age and its associated error. In practice equations [1] and [2] can be replaced by any model and following recent works advocating the use of quantile regression (Van Etten and Hijmans 2010, Silva et al 2015, Riris and Silva 2021) we use an asymmetric Laplace likelihood (Yu and Moyeed 2001), so that $\theta$ is defined as follows:

$$\theta_i \sim \text{AsymLaplace}(\mu_i, \lambda, \tau) \quad [4]$$

where $\mu_i$ is the same linear model as equation [2] above, and $\lambda$ is the scale parameter, and the $\tau$ is the user-defined quantile of interest. Because we are interested in the distribution of the earliest dates, previous applications of quantile regression for dispersal processes employed extreme values such as 90th ($\tau$=0.9) and 99th percentiles ($\tau$=0.99).

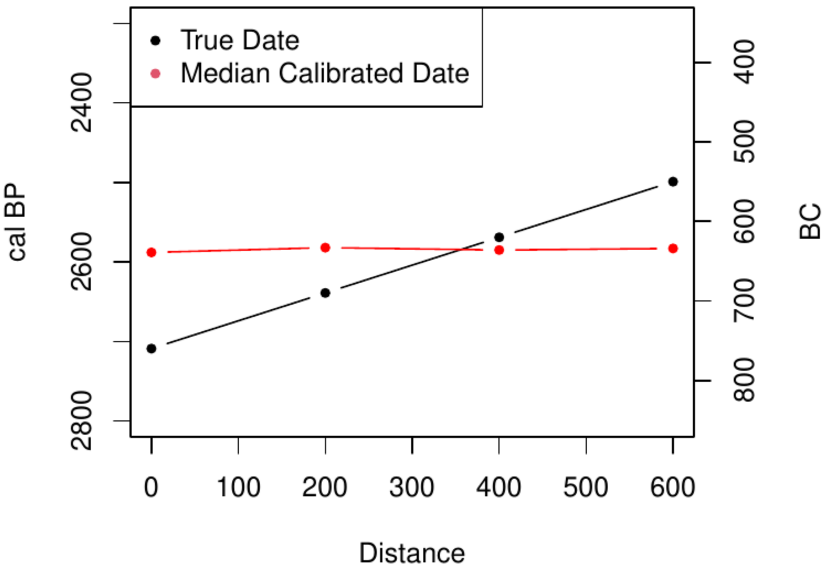


**Fig.S1**. Impact of systematic measurement error on a putative relationship between distance and time for dates falling the calibration plateau. True rate (black) shows a positive slope indicative of dispersal rate of ca 2.85 km/year, whilst median calibrated dates (red) have a flat slope with an instantaneous (i.e. infinite) dispersal rate.

Figure S2 shows the result of the quantile regression model described above fitted to the earliest charred rice dates from each archaeological site, using as the putative source of dispersal the earliest date in our sample (Ukikunden Shell Midden in Northern Kyushu; 130.0087, 33.411, Fig.1 main text). We fitted our model using the following priors:

$$\beta_0 \sim \text{Normal}(3000, 200) \quad [5]$$
$$\beta_1 \sim \text{Exponential}(1) \quad [6]$$
$$\lambda \sim \text{Exponential}(0.01) \quad [7]$$

setting $\tau$ equal to 0.9.

Posterior estimates of the model parameters were obtained using Markov Chain Monte-Carlo (MCMC) sampling via the *nimble* (de Valpine et al 2017) and *nimbleCarbon* (Crema and Shoda 2021, Crema 2021) packages in R (R Core Team 2022). We used four chains, each with 6 million iterations (half dedicated to burn-in) and parameters sampled every 300 steps to reduce file size. We used Metropolis-Hasting adaptive random walk sampler for all parameters using default settings, with the exception of the age estimates $\theta$, where we used an *adaptInterval* of 20,000 and an *adaptFactorExponent* to 0.1 which yielded better mixing in test runs. Convergence was evaluated using the Gelman-Rubin diagnostic and visual inspection of the trace plots. Results showed good mixing with the highest R-hat equal to 1.015..

Figure S2 contrasts this Bayesian model (shown in red) against a non-Bayesian quantile regression model fitted with the median calibrated dates using the *quantreg* (Koenker 2021) R package. The result mirrors the expectations of figure S1, with the median date model (in blue) showing a flatter slope and consequently a much higher expected rate of dispersal compared to the Bayesian model (in red). This discrepancy is due to the impact of the so-called Hallstatt plateau in the calibration curve and is made evident by the larger discrepancies between the median calibrated dates and the median posterior values of $\theta$. The discrepancy is almost absent for periods where the impact of the plateau is small and the model overall provides little additional information concerning individual dates. However, dates in the plateau region, which typically show wider and flatter posterior calibrated distribution are largely "informed" by the overall model, with a resulting shrinking of the measurement error, and in this case an offset of the resulting median date. While it is worth noting that this model does not account for regional variation in the dispersal rates, the reciprocal of the posterior of the parameter $\beta_1$ (figure S3) provides an estimate of the average dispersal rate of charred rice farming between 1.55 and 2.68 km/year.

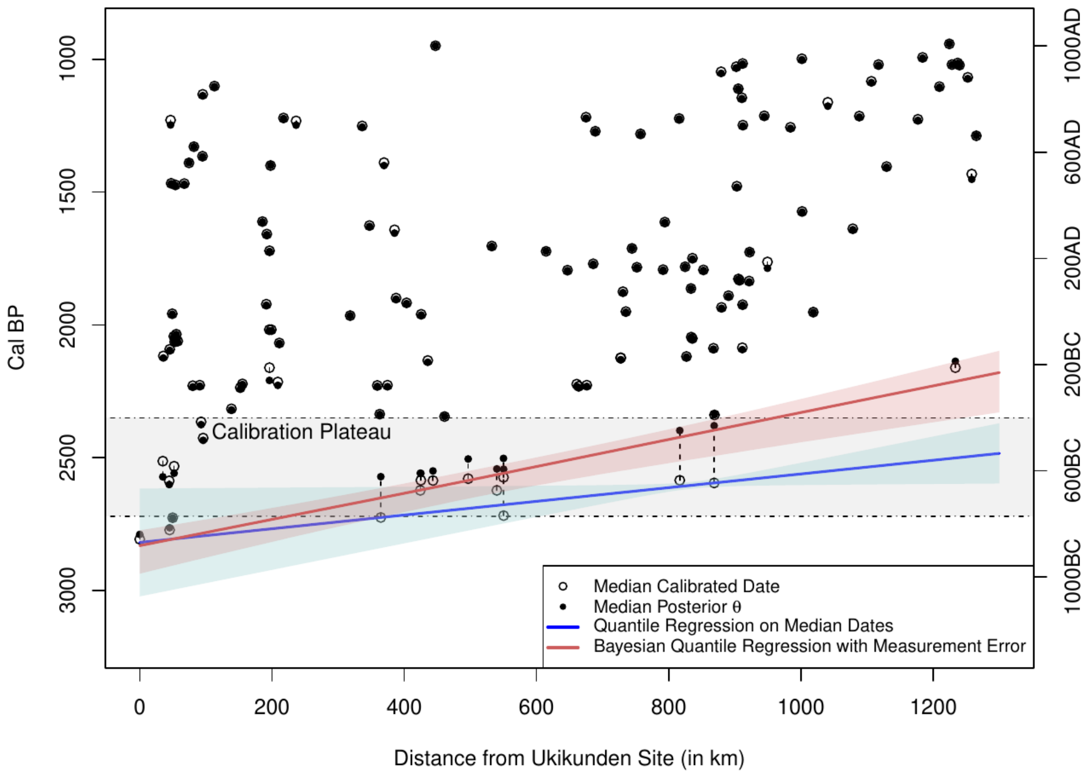

**Fig.S2.** Quantile regression model fitted on median calibrated date (blue line) and accounting for measurement error via Bayesian hierarchical model (red line). Hollow dots are the median calibrated dates while the filled dots are the median posterior of the Bayesian model accounting for the overall model. The offset between median dates and the model posterior are largest in the calibration plateau region, where the calibrated dates generally show flat distributions and the overall model can provide the highest amount of additional information.

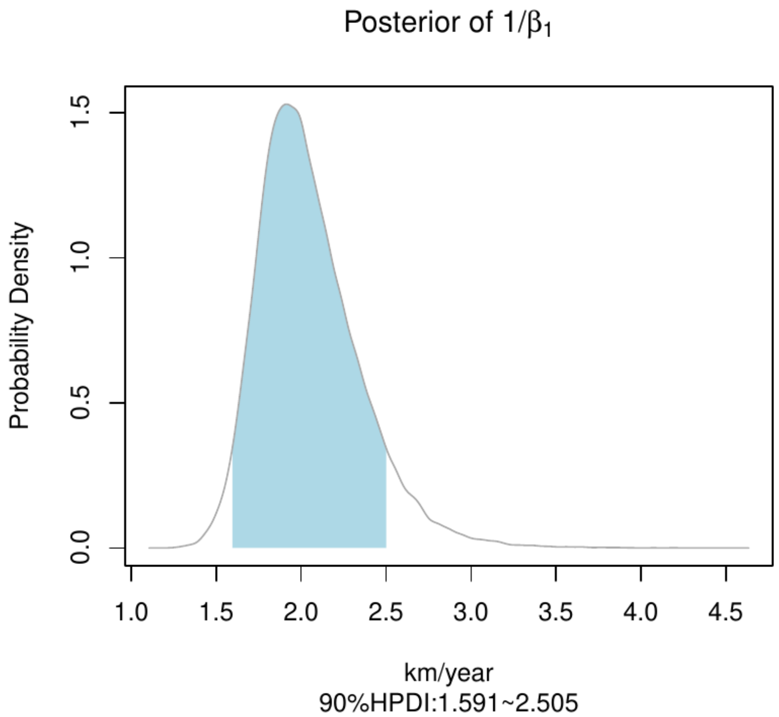


**Fig. S3**. Estimated dispersal rate for the spread of charred rice farming in Japan using Bayesian quantile regression.

### 2. Bayesian Gaussian Process Quantile Regression

The linear model formalised in equation [2] assumes that $-1/\beta_1$ is constant across the geographic window of analyses and hence not able to identify local episodes of acceleration and slow-down in the dispersal process. A slight modification of the linear model can account for such local processes:

$$\mu_i = \beta_0 + d_i (s_i - \beta_1) \qquad [7]$$

Here the additional term $s_i$ allows for a site-level variation in the dispersal rate. This local deviation parameter can be assumed to be a random effect within a hierarchical setting, but with the important additional assumption that $s$ is spatially autocorrelated. A Gaussian process model (Rasmussen and Williams 2016) can satisfy these two assumptions so that the parameter is modelled as a multivariate normal distribution:

$$s_i \sim \text{MVNormal}(0, \mathbf{\Sigma}) \qquad [8]$$

with a mean of 0 and with a covariance matrix $\mathbf{\Sigma}$ defined by a quadratic exponential model (see also figure S4):

$$\mathbf{\Sigma}_{i,j} = \eta^2 \exp(-0.5(d_{i,j}/\rho)^2) + I_{i,j}\varepsilon^2 \qquad [9]$$

where the covariance $\mathbf{\Sigma}_{i,j}$ between sites *i* and *j* is portrayed as an exponential decay based on their inter-distance $d_{i,j}$ with length scale parameter ρ and maximum covariance equal to the square of the marginal deviation parameter η. The term $I_{i,j}\varepsilon^2$ provides an additional covariance $\varepsilon^2$ in case the *i* and *j* are identical.

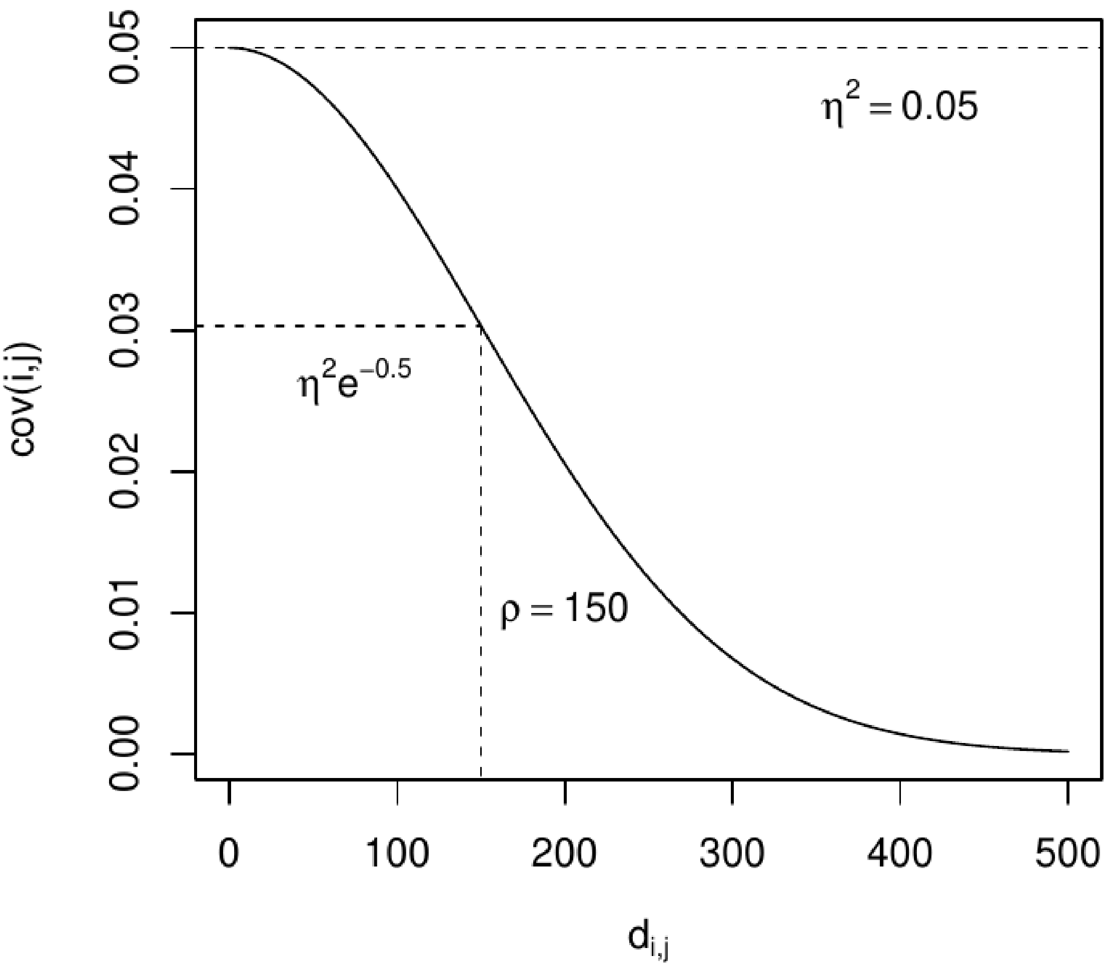


**Fig.S4**. The quadratic exponential model and its relationship to its parameters ρ, η, and $d_{i,j}$ assuming *i≠j*.

Figure S5 shows the result of simulated dispersal rates under different settings of ρ and η. With all things equal, larger values of ρ yield wider regions with similar dispersal rates, whilst larger values of η lead to a wider range of values of *s*, and consequently greater diversity in the dispersal rate.

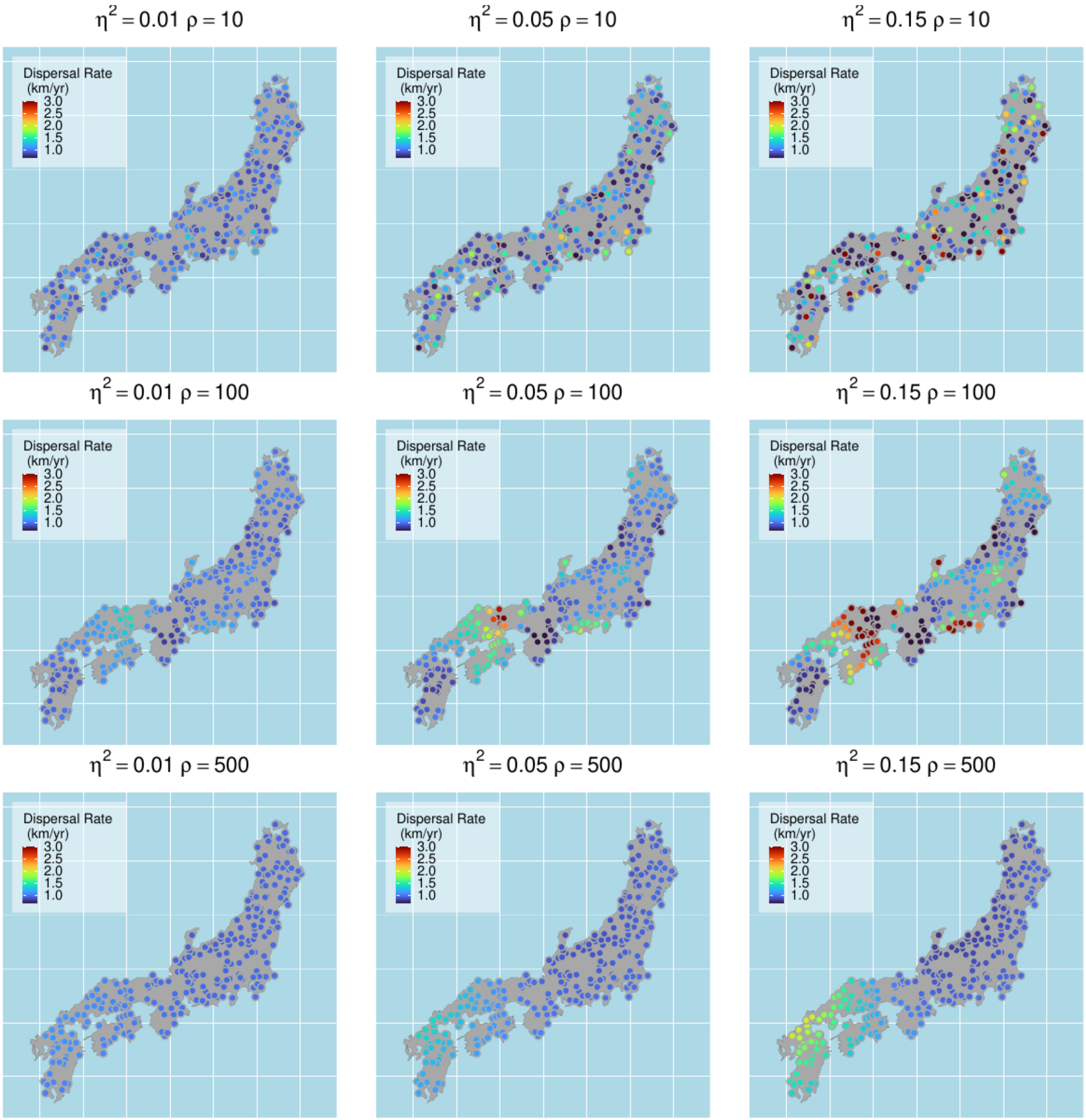


**Fig.S5**. Simulated local dispersal rates with different settings of ρ and η, with 300 site locations and $\beta_0 = 3000$ and $\beta_1 = 1$

It is worth mentioning here that the local deviation parameter *s* is based on the slope and hence should not be directly interpreted in terms of dispersal rate. For example with $\beta_1$ = -0.5 and *s*=0, the dispersal rate is equal to 2 km/year (i.e. -1/(0-0.5)), with *s*=0.2 the rate increases to ca. 3.33km/year (i.e an increase of 1.33 km/year), whilst *s* = -0.2 decreases the rate to 1.42 km/year (i.e. a decrease of 0.58 km/year). In other words, the impact of *s* is symmetric towards the slope, but not towards the rate. This is due to the very nature of the reciprocal of slopes which accelerates quickly towards 0 with a limit of infinite velocity when the slope is equal to 0 (see figure S6).

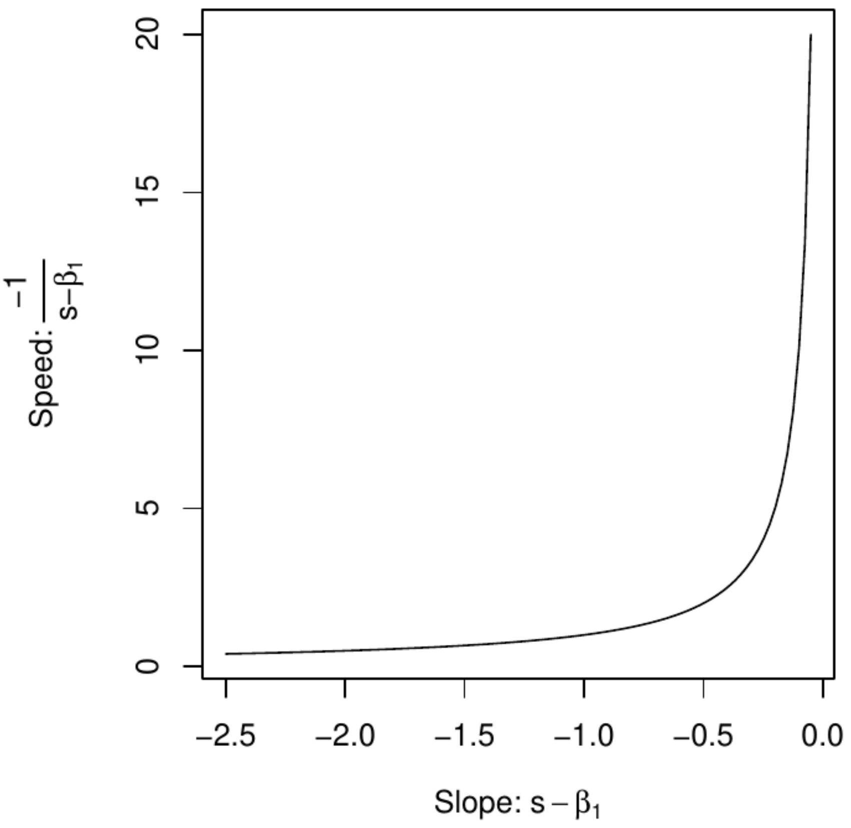


**Fig.S6**. Relationship between regression slope (here $s$-$\beta_1$) and its negative reciprocal, i.e. rate of dispersal.

### 2.1 Priors

Priors for the Gaussian Process Quantile Regression (hereafter GPQR) model were weakly informed by independent lines of archaeological evidence. As for the non-spatial quantile regression model described above, the intercept was based on a normal distribution with a mean of 3,000 and a standard deviation of 200 (see equation [5]) which broadly assumes the arrival of rice to be somewhere between the second half of the 2nd millennium BC and the first half of the 1st millennium BC. Sensible priors for the slope parameter can be based on known archaeological examples of farming dispersal rates, for our case this is equivalent to a reciprocal of $s$ - $\beta_1$ . The variance of the local dispersal rate $s$ is equal to $\eta^2$ in the case of a complete lack of spatial autocorrelation. This can be used as the basis for building a prior predictive check model to establish which prior combination yields sensible dispersal rates. Figure S7 shows the result when an exponential prior with a rate of 1 is applied for $\beta_1$ (cf. equation [6]) and an exponential prior with a rate of 20 is used for $\eta^2$. The results have a wide range of dispersal rates that include observed values from other archaeological contexts. Given the premise and assumptions of the spread of rice farming in Japan, the parameter was also censored to ensure that the dispersal rate is always positive.

The prior for the length scale parameter ρ was modelled using a gamma distribution modelled with a shape of 10 and a rate of 0.06 and truncated between 1 and 1350 (the smallest and largest inter-distance between sampled sites). The gamma distribution enforces positive values, with a fairly long tail and in this case a mode around 200km which ensures a fairly wide range of scales for regional patterns of local dispersal rates (Fig. S8).

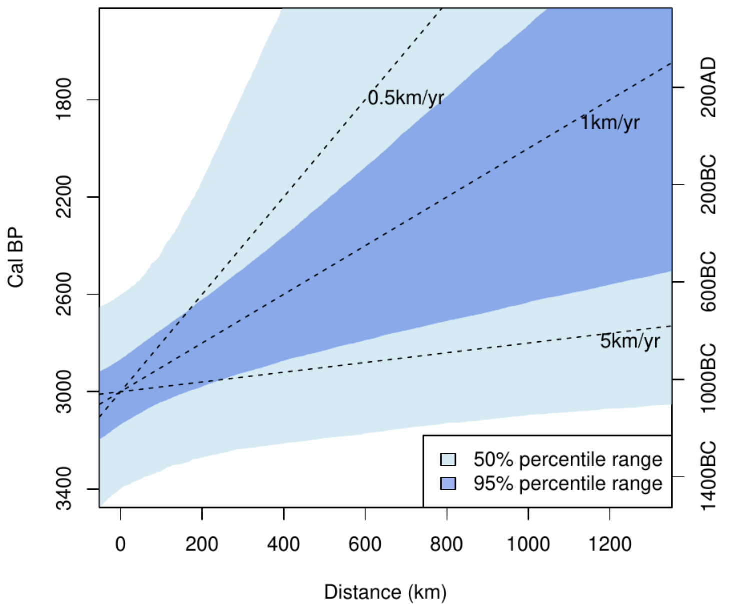


**Fig.S7**. Prior predictive check for $\beta_0$ (modelled as a Gaussian with a mean of 3000 and a standard deviation of 200), $\beta_1$ (modelled as an exponential with a rate of 1), and $\eta^2$ (modelled as an exponential with a rate of 20). The slope was derived by sampled values of s - $\beta_1$, with s drawn from a gaussian with a mean of 0 and a standard deviation equivalent to η. Dashed lines represent the slope for specific rates of dispersal.

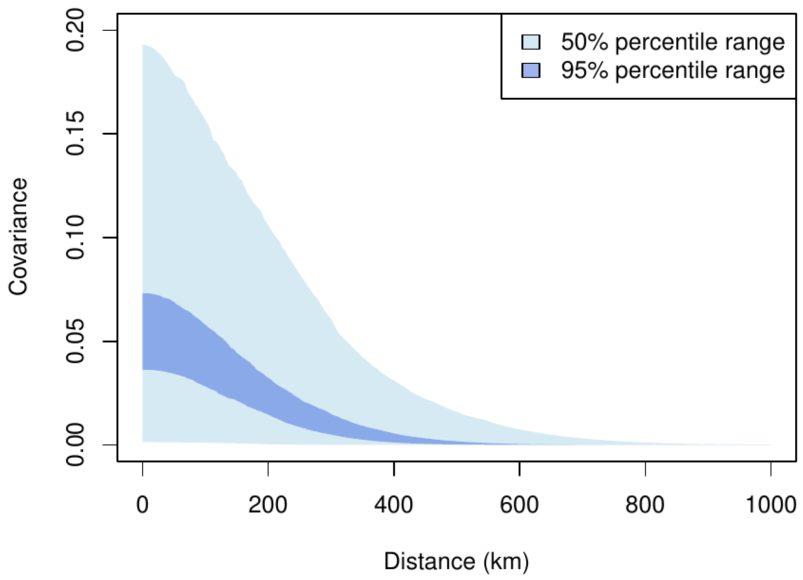


**Fig.S8**. Prior predictive check for $\eta^2$ (modelled as an exponential with a rate of 20) and ρ (modelled as a truncated gamma distribution with a shape of 10 and a rate of 0.06, bounded between 1 and 1350).

### 2.2 Tactical Simulation

To establish whether the Gaussian Process model described above is capable of recovering local variations in dispersal rate we generated a simulated set of $n$=150 radiocarbon dates from an equivalent number of sites within our window of analysis. Calendar dates were sampled using a normal distribution (see equations [1], [7-9] above) using the following parameter values: $\varepsilon = 100$ ; $\beta_0 = 3000$; $\beta_1 = 0.6$; $\rho = 150$; and $\eta^2 = 0.08$. Dates were back-calibrated in $^{14}C$ age with an error of 20 years.

We fitted our dataset using our Gaussian Process model using the same prior as in our observed dataset and setting $\tau$ to 0.9. Model fitting was performed using four chains with 1 million iterations each (half discarded as burn-in, and sampled every 50 steps). MCMC diagnostics showed good convergence, with the highest R-hat (Gelman and Rubin 1992) equal to 1.003.

Our results indicate that the model does capture the spatial variation in dispersal rates (figure S9), with most of the random effect parameters (figure S10) and all the key fixed effect parameters (figure S11) recovering their respective true values within their 95% quantile intervals.

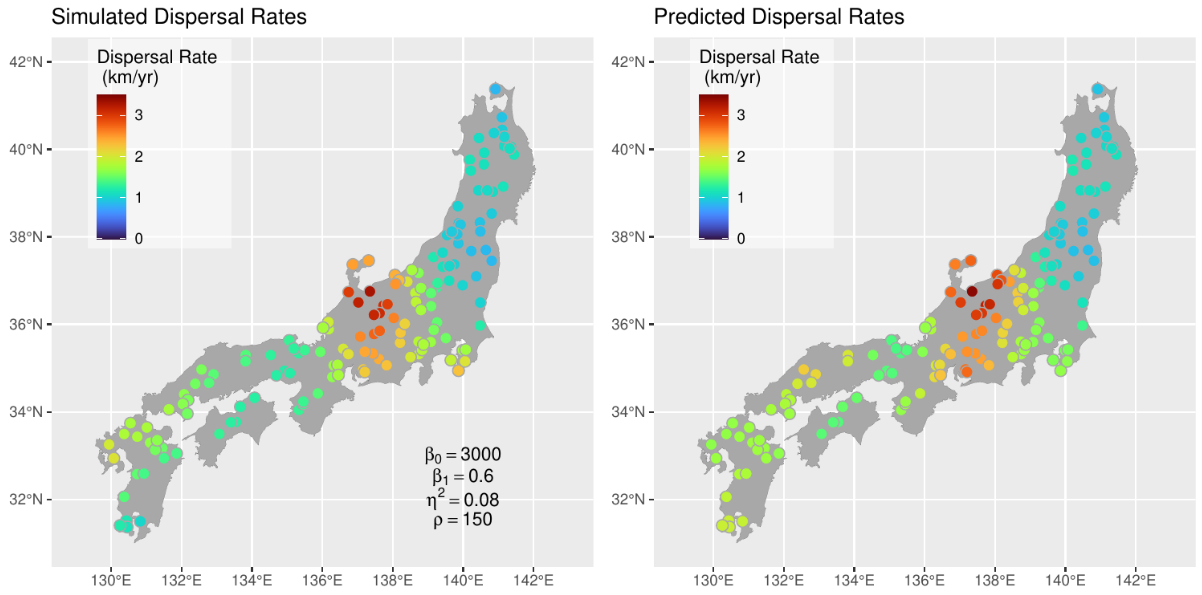


**Fig.S9**. Simulated and predicted (median posterior) dispersal rates.

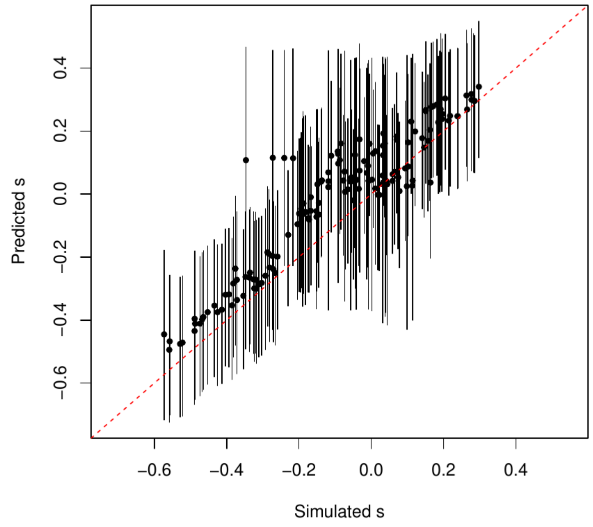


**Fig.S10**. Simulated and predicted values of the spatial random effect parameter *s*.

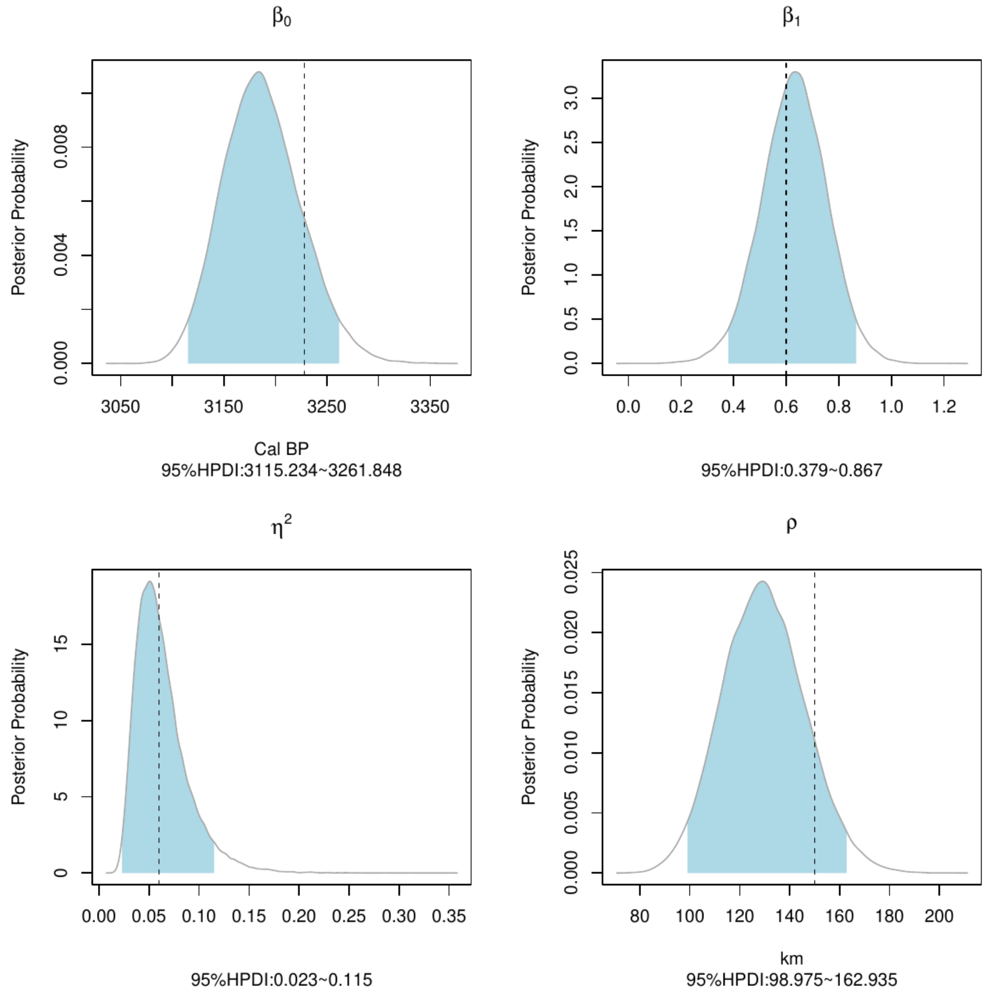


**Fig.S11**. Posterior distribution of key model parameters for the simulated dataset with corresponding "true" values shown as a dashed line. Notice that in the case of $\beta_0$ the line represents the 90th percentile matching the value of $\tau$used for inference.

### 2.3 MCMC Settings, Diagnostics, and Posteriors

As for the non-spatial quantile regression the model fitting was carried out using the *nimble* and the *nimbleCarbon* R packages. We ran four chains, each with 2 million iterations, half of which were used for burn-in, and values sampled every 100 steps. We used a Metropolis-Hasting adaptive random walk sampler for all parameters with the exception of $\beta_1$ and $s_i$ in the GPQR model, which were inferred using an automated factor slice sampler (Tibbits et al 2014) to account for correlation between the two parameters. As for the non-spatial quantile regression, the random walker sampler for $\theta$ was modified from the default settings (see section 1 above).

The Gelman-Rubin convergence statistics (Gelman and Rubin 1992) were below 1.01 for all parameters, and the trace plots of the four key fixed parameters showed good mixing of the chains (fig. S12 & S13). Figures S14 and S15 show the marginal distribution of these four parameters, whilst tables S1 and S2 provide summary statistics and diagnostics of all fixed parameters.

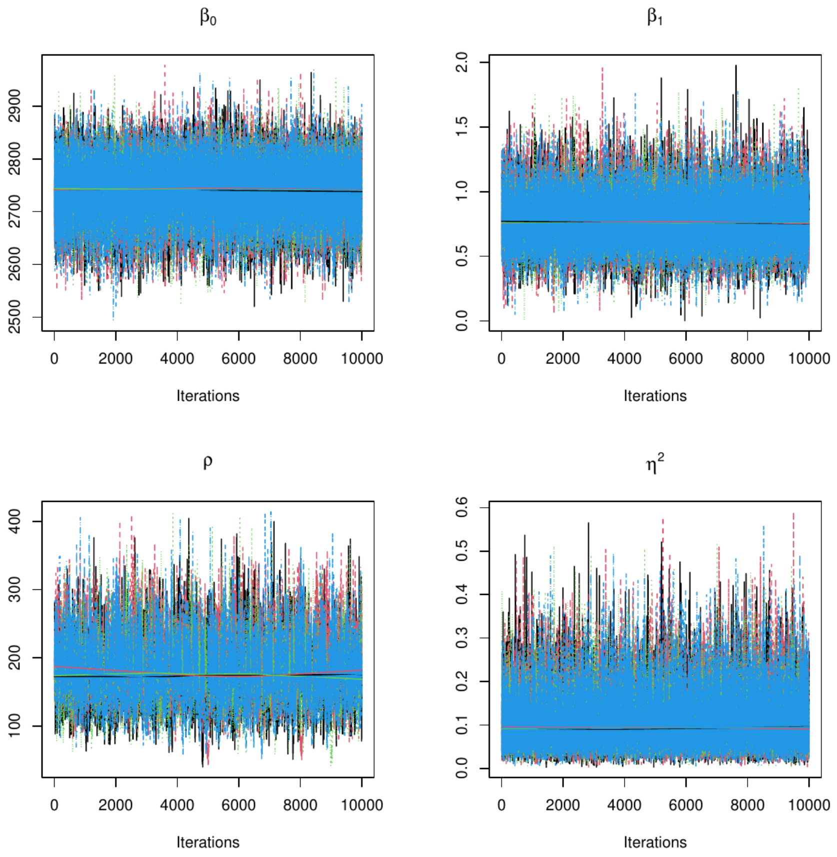


**Fig.S12**. Traceplots for $\beta_0$, $\beta_1$, ρ, and $\eta^2$ for the $\tau$=0.9 GPQR model.

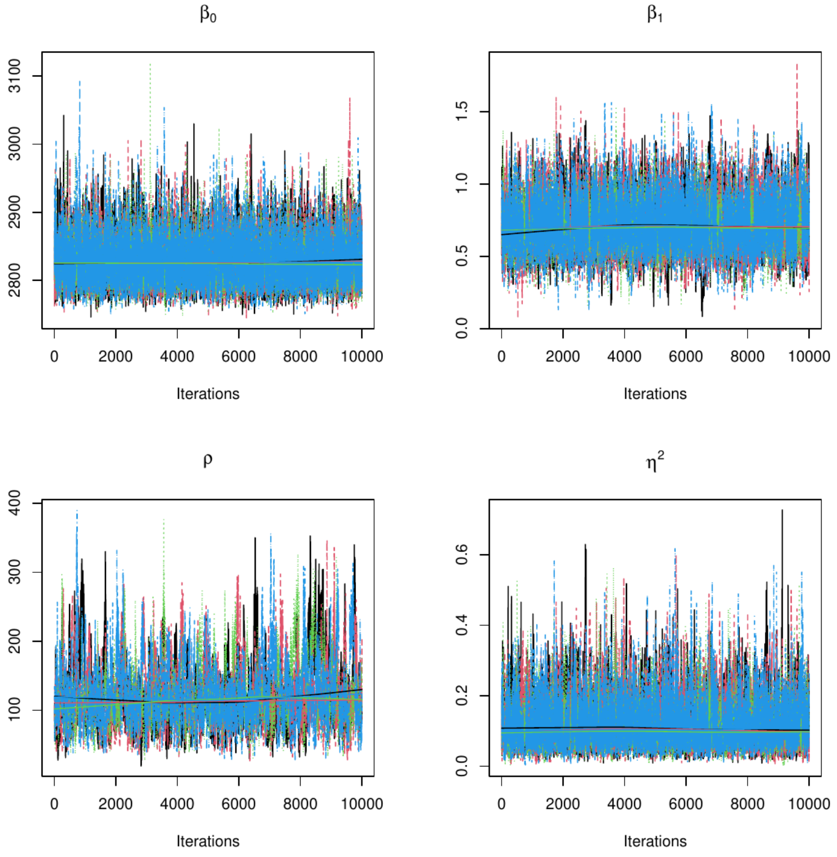


**Fig.S13**. Trace Plots for $\beta_0$, $\beta_1$, ρ, and $\eta^2$ for the $\tau$=0.99 GPQR model.

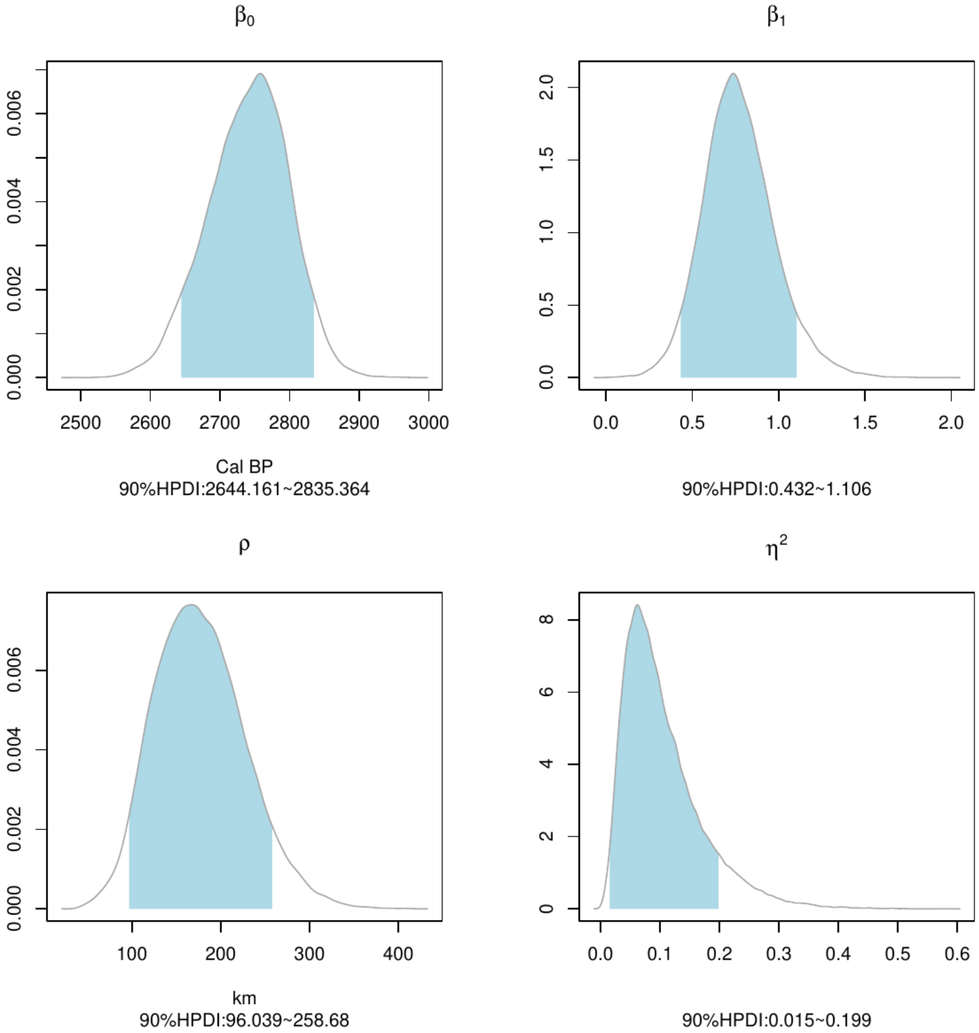


**Fig.S14**. Marginal posterior distribution of $\beta_0$, $\beta_1$, ρ, and $\eta^2$ in the $\tau$=0.90 GPQR model.

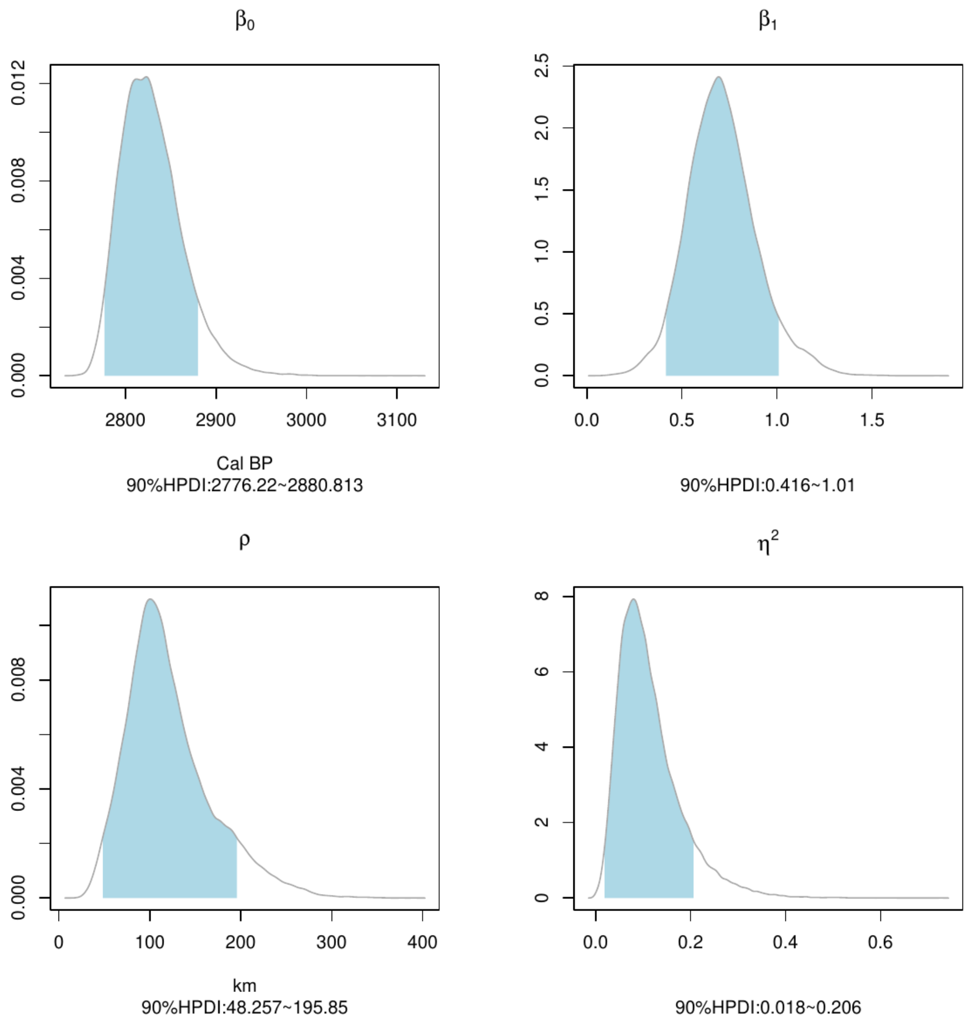


**Fig.S15**. Marginal posterior distribution of $\beta_0$, $\beta_1$, ρ, and $\eta^2$ in the $\tau$=0.99 GPQR model.

| Parameter | Median Posterior | 90% HPDI (low) | 90% HPDI (high) | R-hat | ESS |
|---|---|---|---|---|---|
| $\beta_0$ | 2744 | 2644 | 2835 | 1.00 | 22266 |
| $\beta_1$ | 0.75 | 0.43 | 1.10 | 1.00 | 28394 |
| ρ | 173.70 | 96.03 | 258.68 | 1.00 | 3032 |
| $\eta^2$ | 0.0888 | 0.0147 | 0.1992 | 1.00 | 23224 |

**Table S1** Summary statistics and diagnostics for $\beta_0$, $\beta_1$, ρ, and $\eta^2$ in the $\tau$=0.90 GPQR model.

| Parameter | Median Posterior | 90% HPDI (low) | 90% HPDI (high) | R-hat | ESS |
|---|---|---|---|---|---|
| $\beta_0$ | 2825 | 2776. | 2881 | 1.00 | 7432 |
| $\beta_1$ | 0.70 | 0.42 | 1.01 | 1.00 | 2704 |
| ρ | 112.32 | 48.26 | 195.85 | 1.01 | 765 |
| $\eta^2$ | 0.0998 | 0.0185 | 0.2063 | 1.01 | 3475 |

**Table S2** Summary statistics and diagnostics for $\beta_0$, $\beta_1$, ρ, and $\eta^2$ in the $\tau$=0.99 GPQR model.

### 3. Bayesian Hierarchical Phase Models with Constraints

Bayesian phase models of radiocarbon dates typically used in stratigraphic contexts have been occasionally applied to study regional phenomena, such as the dating of cultural phases (Ziedler et al 1998, Brunner et al 2020, Crema and Kobayashi 2020) or the arrival date of populations (Blackwell and Buck 2003) or cultural traits (Capuzzo and Barceló 2015). These models have been recently used to estimate the arrival dates of crops in different geographic regions (Long et al 2020, Leipe et al 2020), typically employing uniform distribution models as follows:

$$\theta_{i,k} \sim \text{Uniform}(\nu_k, \upsilon_k) \qquad [10]$$

where $\theta_{i,k}$ is the calendar date of sample $i$ from Area $k$, and $\nu_k$ and $\upsilon_k$ are the start and end dates of the Area $k$. The relationship between $\theta_{i,k}$ and the radiocarbon sample is established using fundamentally the same procedure described above in the model [2].

The parameter of interest, in this case, is $\nu_k$ which provides an estimate of the arrival time of the crop in the focal Area $k$. While there are some interesting non-Bayesian alternatives to this problem (Key et al 2021), the advantage of this approach is the possibility to add constraints and priors in a multi-regional setting (see below).

A key assumption of this approach, however, is that in order to have unbiased estimates of $\nu_k$ and $\upsilon_k$ samples must be randomly and independently selected. For many real statistical applications, this assumption is never truly met, but in the case of crop dispersal, a particular problem arises from the fact that charred remains are often found in distinctive recovery contexts or sites, and as such samples cannot be regarded as truly independent. A trivial example is to contrast a situation where the inference is carried out on 20 samples from 20 different sites against a situation where the same number of samples are recovered from a single site. In the former case estimates of $\nu_k$ and $\upsilon_k$ are more representative of the start and end dates of a region, but in the latter case, these would represent the start and end date of the particular site from which the samples are taken. While such extreme situations are rare, multiple samples are frequently taken from the same site or context, and as such might represent a substantial portion of the dates from a particular region. For example, Leipe et al 2020 estimated the arrival date of rice farming in the Japanese central highlands using 32 dates, but ca. 40% of them (n=13) were from a single site (Chikarishijori).

One way to approach this problem and to take into account the fact that samples are typically recovered from the same site is to employ a hierarchical model:

$$\theta_{i,j,k} \sim \text{Uniform}(\alpha_{j,k}, \alpha_{j,k} + \delta_{j,k}) \qquad [11]$$
$$\alpha_{j,k} \sim \text{Uniform}(\nu_k, \upsilon_k) \qquad [12]$$

where $\alpha_{j,k}$ is the start date of crop use at site $j$ in region $k$, and $\delta_{j,k}$ is the duration of rice farming in the same site. Inference of the regional start date is then no longer based on the individual charred dates $\theta_{i,j,k}$ but instead from the start date of individual sites. This effectively acknowledges sample independence; sites with a larger number of dates will still provide more information, but mainly in terms of having more accurate estimates of $\alpha_{j,k}$. From an interpretative point, [12] is slightly different from [10] as the former represents the distribution of start dates at each site while the latter represents just the distribution of dates within the area $k$. However, the interpretation of our primary parameter of interest (i.e. $\nu_k$) as the arrival time of the crop in the partictical area remains the same. Model [11] is parameterised in a way so that the end date is effectively defined by the duration parameter $\delta_{j,k}$. Inference of this parameter is actually helped by the presence of sites yielding multiple dates and by the choice of some weakly informative prior that conveys assumptions on site duration. Of course, if the entirety of the dataset consists of one date per site then there is effectively no way to infer $\delta_{j,k}$, but under those circumstances arrival dates can be safely estimated using model [10].

#### 3.1 Tactical Simulation

To illustrate the difference between the standard model [10] and the proposed hierarchical modification depicted in [11] and [12], we simulated an artificial archaeological dataset consisting of 30 radiocarbon dates distributed exponentially across 10 archaeological sites within a single region ($n_1 = 13$, $n_2 = 3$, $n_3 = 5$, $n_4 = 3$, $n_5 = n_6 = n_7 = n_8 = n_9 = n_{10} = 1$). Start dates of occupation (i.e. $\alpha_j$) were modelled using a uniform distribution with regional start and end dates $\nu$ and $\upsilon$ dates equal to 3500 and 3000 cal BP. The site duration parameter $\delta_j$ was randomly sampled from a gamma distribution with shape parameter 5 and rate parameter 0.02. The simulated calendar dates were back-calibrated in $^{14}C$ age and assigned to an error of 20 years.

We estimated $\nu$ and $\upsilon$ using both the conventional non-hierarchical approach (model [10]) and the proposed solution (model [11-12]) using three chains and 2 million iterations (half discarded for burnin and sampled every 100 steps). For model [10] we used a uniform prior for $\nu$ and $\upsilon$ between 0 and 10000, with the constraint $\nu > \upsilon$ (i.e. $\nu$ earlier than $\upsilon$). For model [11-12] we additionally modelled $\delta_j$ using a uniform hyper-prior for the rate (bounded between 1 and 20) and a normal distribution with a mean 200 and standard deviation 100 truncated between 1 and 500 for the mode, where the rate is equal to (shape-1)/mode. MCMC diagnostic showed good convergence for all parameters (Rhat < 1.01).

Figure S16 shows the difference in the marginal posterior distribution of $\nu$ and $\upsilon$ in the two models. The conventional model has a narrower posterior given the inference is made on 30 samples, whilst the hierarchical model has a wider posterior as the inference is primarily driven on the sites. However, the posterior of the conventional model is less accurate and fails to capture the true start date (dashed line), which is instead recovered by the hierarchical model.

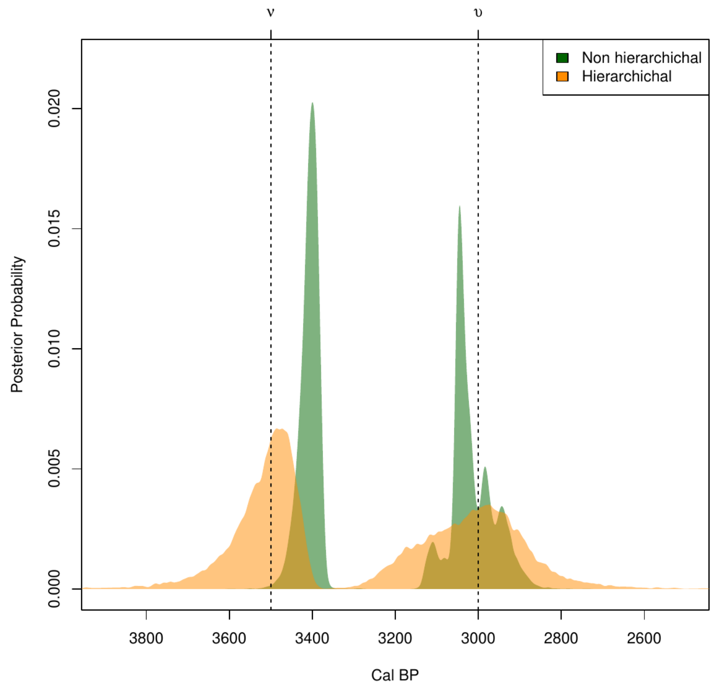


**Fig.S16.** Marginal posterior distribution of the stard and end dates ν and υ in the hierarchical and non-hierarchical models .

### 3.2 Implementation details for the observed data

The hierarchical phase model described in section 3.1 was fitted to the charred rice dates from the eight geographic areas (table S3), whose boundaries were informed by previous archaeological studies and the results of our GPQR analysis, to ensure that dispersal rates within each area were as consistent as possible. We particularly made an effort to avoid having instances where a major change in dispersal rate occurred within each area, so that an early estimate of arrival date was the consequence of dates occurring at sites located at the boundary of adjacent areas.

| Area | Prefectures | Number of Dates | Number of Sites |
|---|---|---|---|
| I | Saga \| Fukuoka \| Nagasaki | 32 | 19 |
| II | Miyazaki \| Oita \| Kumamoto \|Kagoshima | 49 | 22 |
| III | Ehime \| Kochi \| Hiroshima \| Okayama \| Tottori \| Shimane \| Yamaguchi | 22 | 14 |
| IV | Hyogo \| Kagawa \| Kyoto \| Mie \| Nara \| Osaka \| Shiga \| Tokushima \| Wakayama | 32 | 11 |
| V | Aichi \| Gifu \| Ishikawa \| Fukui \| Kanagawa \| Niigata \| Nagano \| Toyama \| Shizuoka \| Yamanashi | 94 | 35 |
| VI | Chiba \| Gunma \| Ibaraki \| Tokyo \| Saitama \| Tochigi | 38 | 14 |
| VII | Akita \| Fukushima \| Iwate \| Miyagi \| Yamagata | 18 | 9 |

| VIII | Aomori | 9 | 8 |
|---|---|---|---|
| TOTAL | | 294 | 132 |

**Table S3.** Number of sites and radiocarbon dates from each area

Priors of the start and end dates ν and υ were based on a uniform distribution bounded between 500 and 5000 cal BP, with the constraint ν > υ. The duration parameter δ was modelled using a gamma distribution, with the shape and rate parameters modelled using a uniform hyper-prior for the rate (bounded between 1 and 20) and a normal distribution with a mean 200 and standard deviation 100 truncated between 1 and 500 for the mode, where the rate is equal to (shape-1)/mode. Figure S17 shows the prior predictive check with these settings and illustrates how our model includes a wide range of possible settlement durations.

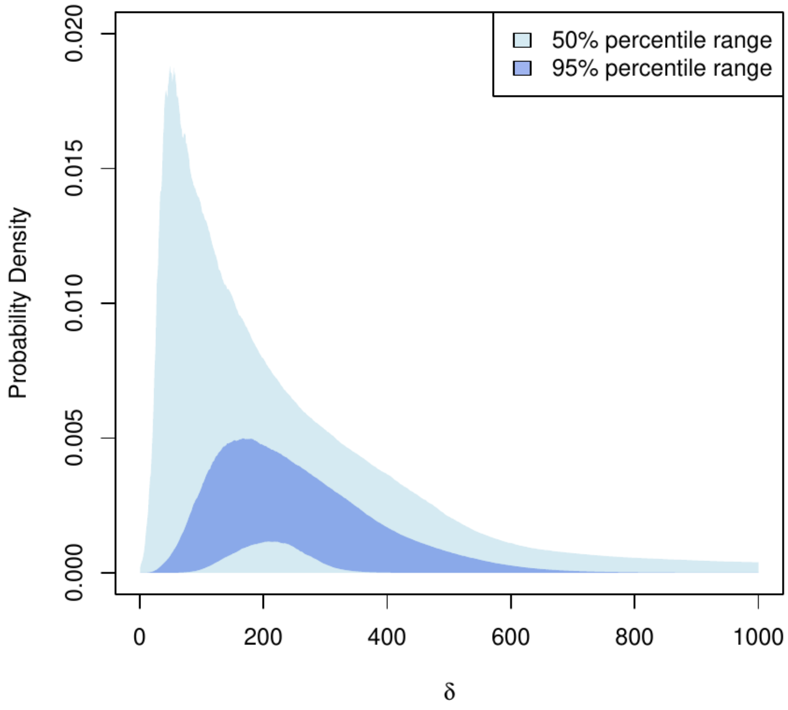


**Fig.S17.** Prior predictive check for the duration parameter δ. Generating using 5,000 random samples from relevant priors.

We considered two different versions of the hierarchical model. In **model a**, we infer all parameters without any constraints, hence allowing for all possible temporal orders of our primary parameter of interest ν. This allows us to infer the arrival date of rice in each area on the exclusive basis of the dates from that focal area. In **model b**, we add the following constraints: $\nu_I > \nu_{kII}$ , $\nu_I > \nu_{III} > \nu_{IV} > \nu_V$ , $\nu_{IV} > \nu_{VI}$ , $\nu_{IV} > \nu_{VII}$ , and $\nu_{IV} > \nu_{VIII}$. This effectively incorporates the assumption of a wave-of-advance like model originating from northern Kyushu (Area I), whilst allowing for possible leap-frogging after the dispersal reached the Kansai area (Area IV).

Both models were fitted using 6 million iterations, half of which were dedicated to burn-in, and sampled every 300 steps. We used Metropolis-Hasting adaptive random walk sampler for all parameters using default settings, with the exception of the age estimates $\theta$, where we used an *adaptInterval* of 20,000 and an *adaptFactorExponent* to 0.1. Model convergence was assessed via visual inspections of the trace plots and Gelman-Rubin convergence statistics. All parameters yielded an R-hat below 1.01, with the exception of one $\theta_{166}$ which returned an Rhat of 1.03 in **model b.** Visual inspection of the trace plot indicated that this was most likely due to the multi-modal shape of the posterior, and the high agreement index (98.4, Bronk-Ramsey 1995), as well as comparison with the associated calibrated date, suggests that there are no concerns from an inferential point of view.

### 3.2 Posterior Distributions

Table S4 shows the summary statistics of the arrival date estimates ν for the two models, whilst figures S18 and S19 show their marginal posterior distribution. Figure S20 indicates the probability that ν values of row areas are smaller (i.e. later) than column areas, with cells coloured in blue supporting a wave-of-advance scenario, while those coloured in red indicate possible leap-frog dispersals. Finally, figure S21 shows the pairwise difference in ν (column area minus row area) obtained from the posterior samples, with portions of the 90% highest posterior density interval coloured in blue for positive values (column earlier than row) and in red for negative values (row earlier than column).

| Model | Area | Median Posterior | 90% HPDI (low) | 90% HPDI (high) | Rhat | ESS |
|---|---|---|---|---|---|---|
| *Model a* | I | 986 BC | 1176 BC | 845 BC | 1 | 39981 |
| | II | 572 BC | 740 BC | 430 BC | 1 | 39424 |
| | III | 887 BC | 1176 BC | 669 BC | 1.001 | 11796 |
| | IV | 905 BC | 1179 BC | 695 BC | 1 | 30453 |
| | V | 655 BC | 771 BC | 552 BC | 1 | 38439 |
| | VI | 273 BC | 478 BC | 122 BC | 1 | 39026 |
| | VII | 153 BC | 458 BC | 44 AD | 1 | 40000 |
| | VIII | 469 BC | 943 BC | 177 BC | 1 | 38182 |
| *Model b* | I | 1039 BC | 1251 BC | 872 BC | 1 | 39658 |
| | II | 570 BC | 735 BC | 430 BC | 1 | 40326 |
| | III | 910 BC | 1061 BC | 779 BC | 1 | 30742 |
| | IV | 824 BC | 946 BC | 703 BC | 1 | 23067 |
| | V | 648 BC | 754 BC | 560 BC | 1 | 36917 |
| | VI | 271 BC | 471 BC | 124 BC | 1 | 35089 |
| | VII | 152 BC | 434 BC | 42 AD | 1 | 39185 |
| | VIII | 428 BC | 709 BC | 203 BC | 1 | 37394 |

**Table S4.** Summary statistics and diagnostics for the ν parameters in the two models.

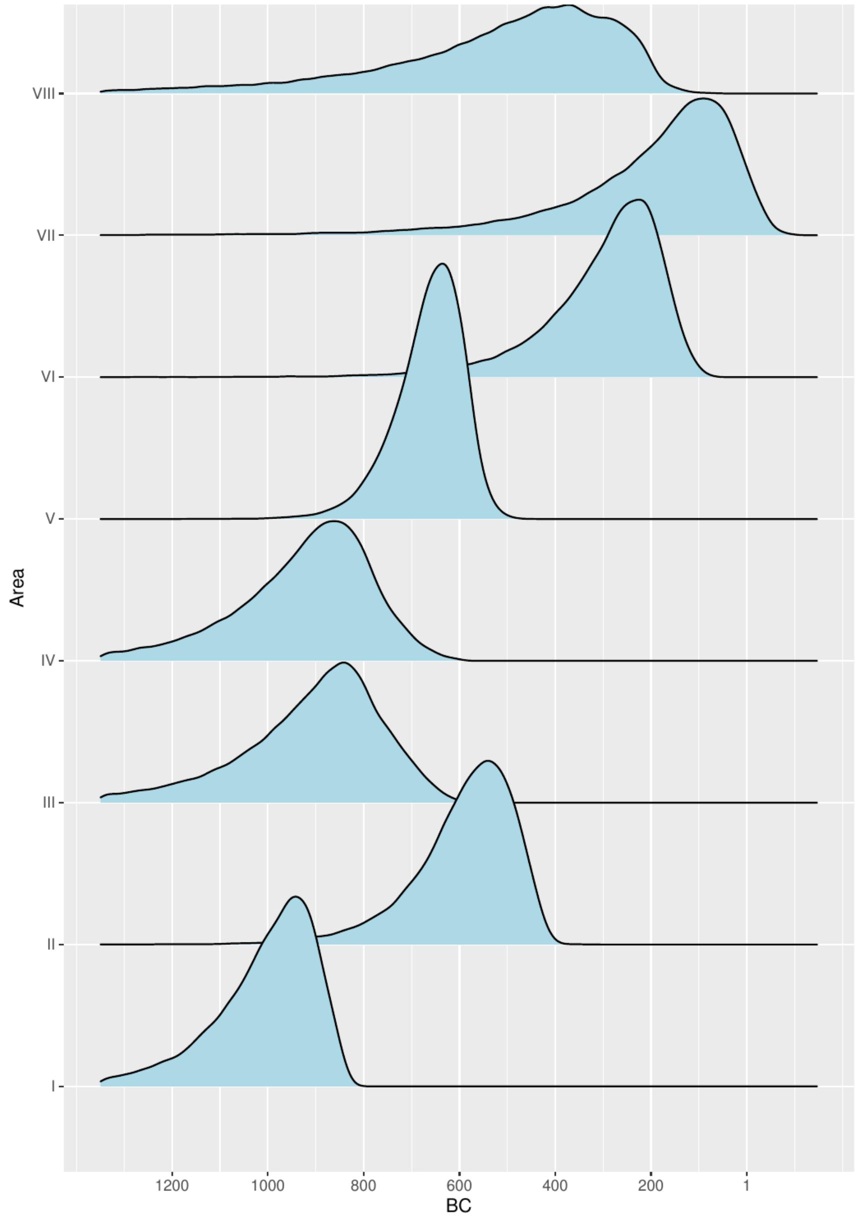


**Fig.S18.** Marginal posterior distribution of ν for model a

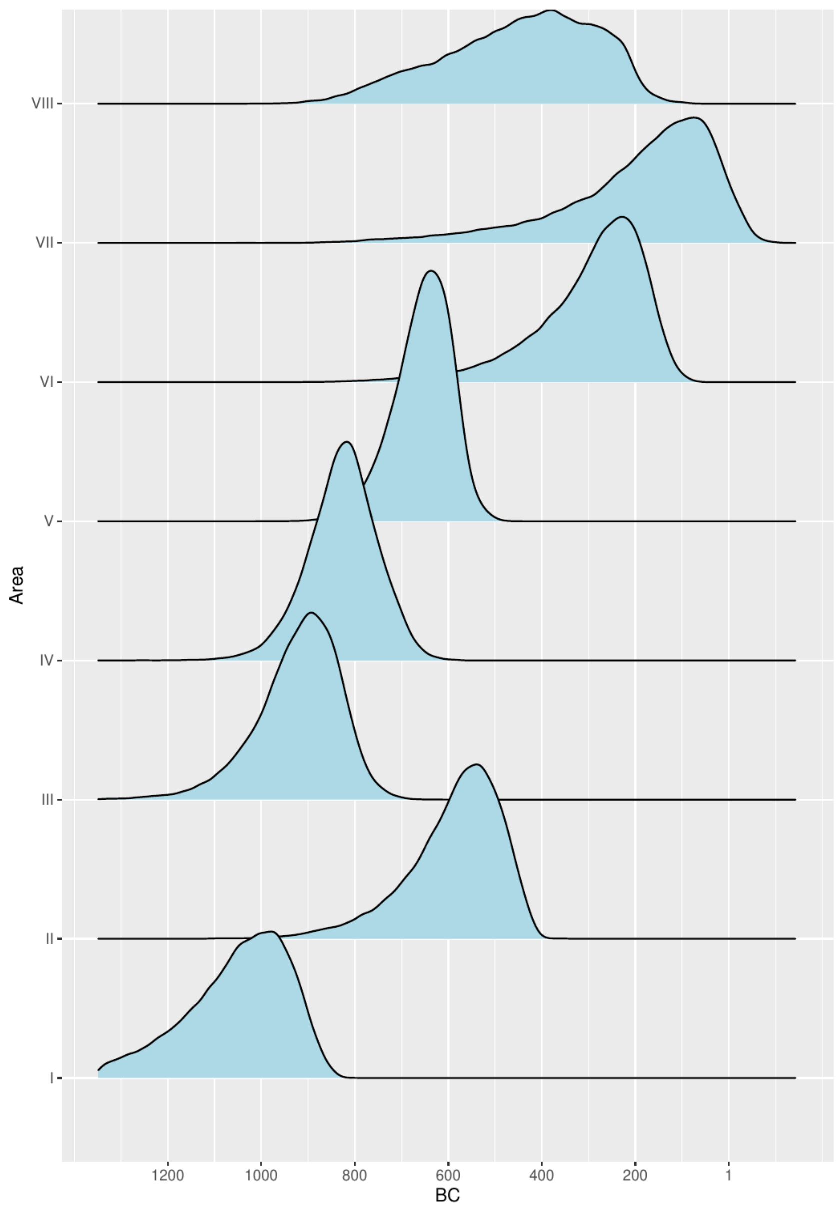


**Fig.S19.** Marginal posterior distribution of ν for model b

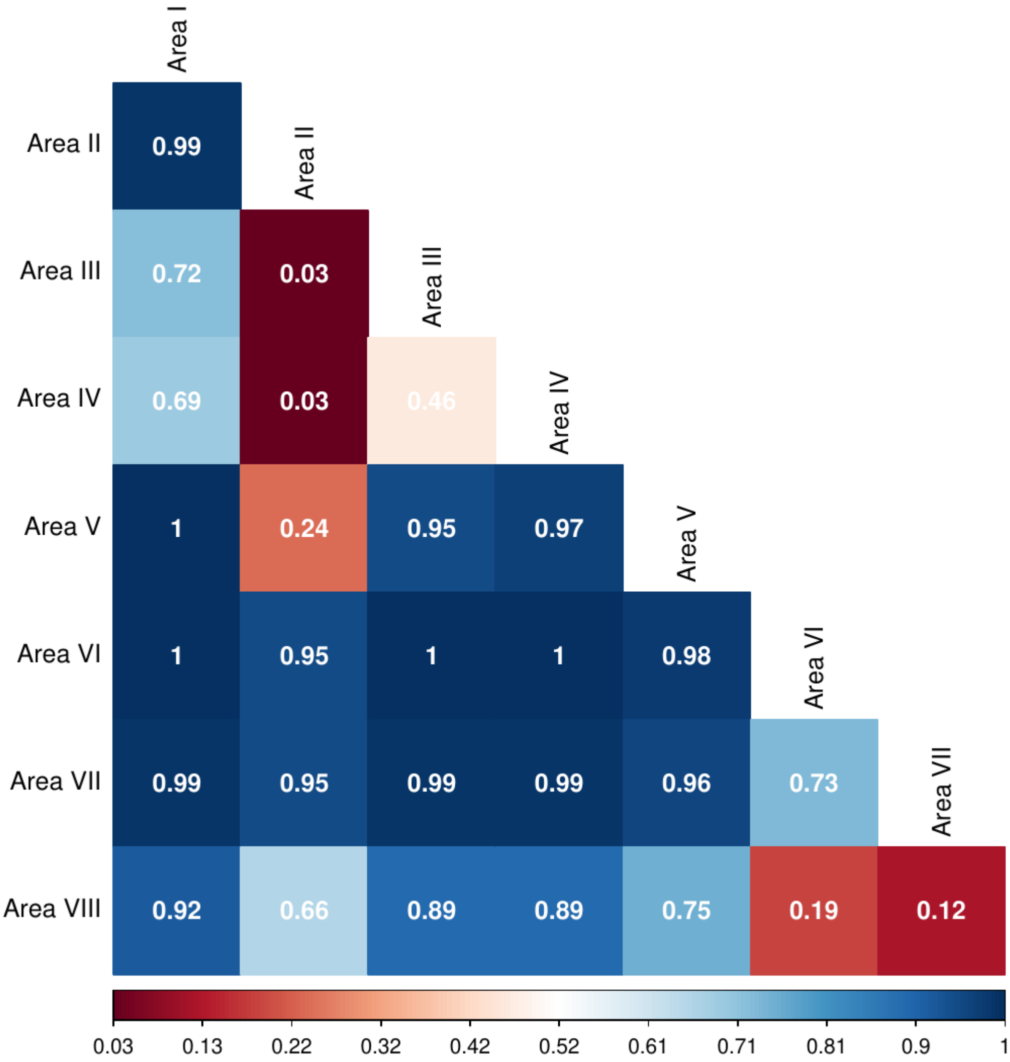


**Fig.S20.** Probability of row areas's v being after colum area's v for model a

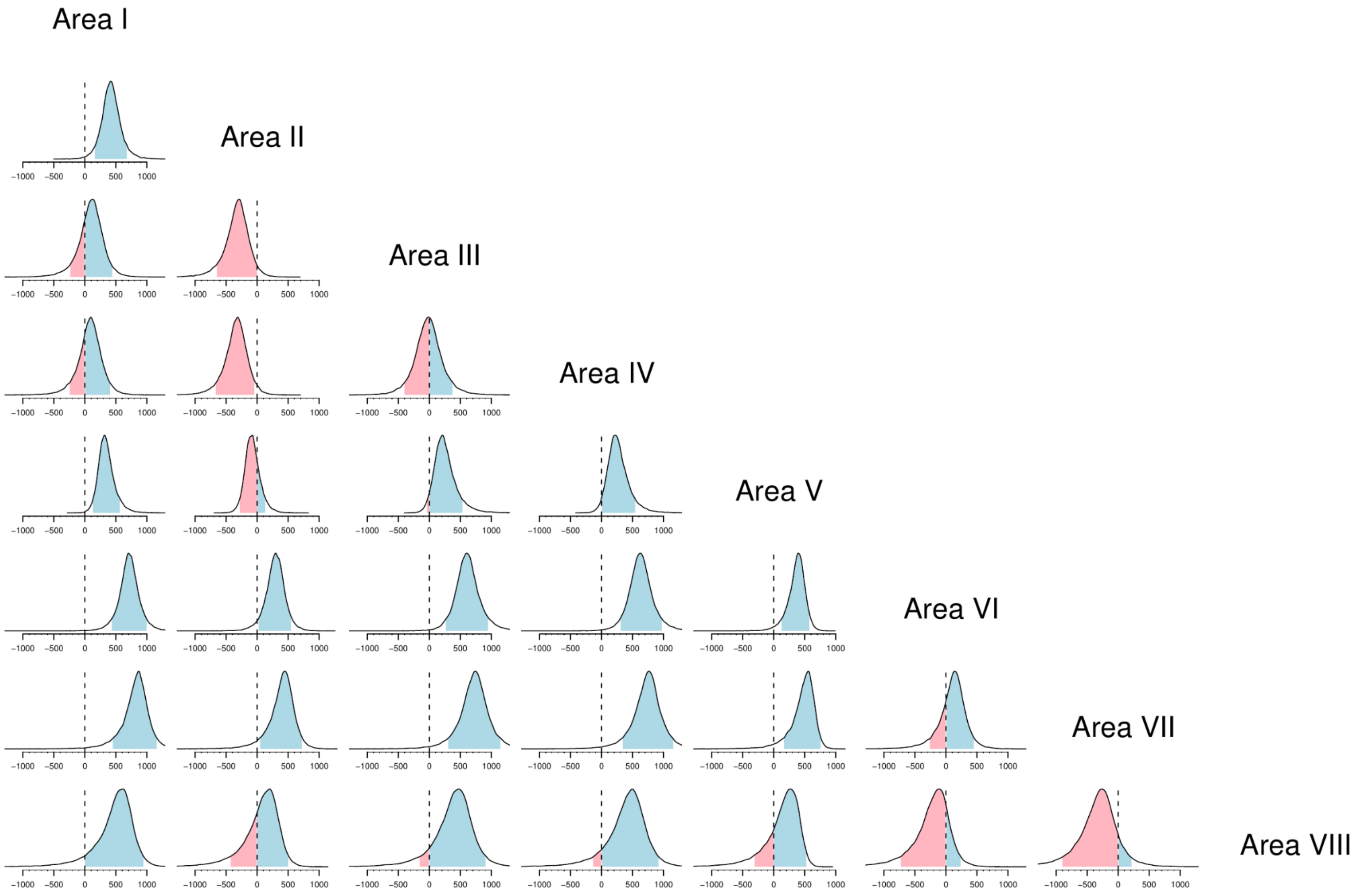


**Fig.S21.** Posterior estimates of pairwise difference (column minus row) of v in model a. Highlighted regions represent the 90% HPDI (red: row is earlier; blue: row is later)
